# Unique nasal cell states induced by common pediatric respiratory viruses

**DOI:** 10.64898/2026.04.20.719671

**Authors:** Jaclyn M. L. Walsh, Lillian J. Juttukonda, Ying Tang, Ankita Chatterjee, Marc Elosua-Bayes, Erica Langan, Amanda Frischmann, Faith Taliaferro, Hannah R. Matthews, Kyle Kimler, Chris M. Lent, Deb Purna Keya, Preonath Chondrow Dev, Apurba Rajib Malaker, Lubana Tanvia, Arif Mohammad Tanmoy, Supratim Ghosh, Arindam Maitra, Apurba Ghosh, Surupa Basu, Abigail S. Kane, Charles A. Coomer, Alex K. Shalek, Thushan I. de Silva, Abdul Karim Sesay, Jessica Edwards, Corrine Sin Quee, Wanda I. Gonzalez, Lael M. Yonker, Sarah C. Glover, Partha Majumder, Yogesh Hooda, Senjuti Saha, Jose Ordovas-Montanes, Bruce H. Horwitz

## Abstract

Respiratory viral infections in early childhood are major drivers of acute morbidity and long-term airway disease, yet how distinct viruses remodel the pediatric nasal mucosa at cellular resolution remains unresolved. Here, we generated a single-cell RNA sequencing atlas of 335,174 nasal epithelial and immune cells from 132 children under five years of age with SARS-CoV-2, rhinovirus, or respiratory syncytial virus (RSV) infection, alongside uninfected controls. Mapping viral transcripts to individual cells revealed virus-specific infected epithelial states: an NF-kB-responsive ciliated subset in SARS-CoV-2 and a previously undescribed *KRT17+* squamous-like subset in RSV. We delineated divergent mucosal response programs, including a robust interferon (IFN) response in SARS-CoV-2, an IL-13-responsive secretory program in rhinovirus, and heightened inflammatory and cytotoxic immune activation in RSV. In RSV, specific immune subsets and elevated IFN-response signatures were associated with disease severity, whereas rhinovirus-induced wheeze was marked by expansion of a *CST1*+ goblet cell subset. Integration of asthma genome-wide association data with our atlas revealed a *KRT13+* hillock-like squamous epithelial subset enriched for expression of childhood-onset asthma risk loci. Finally, we demonstrate that this resource enables high-resolution annotation of independent pediatric cohorts in Kolkata, India and rural Bangladesh. Together, this atlas establishes a comprehensive view of antiviral immunity in the pediatric nasal mucosa and defines virus-specific mucosal immune programs relevant to disease severity and asthma risk in early life.

## Introduction

Respiratory viral infections are exceedingly common in early childhood and represent a major cause of morbidity and hospitalization worldwide. Among these pathogens, respiratory syncytial virus (RSV) and rhinovirus account for a substantial fraction of hospitalizations, wheezing symptoms, and lower respiratory tract disease in infants and young children. RSV alone is responsible for millions of hospital admissions globally each year in children under five^1–5^. In addition to their acute clinical impact, respiratory viral infections in infancy and early childhood have been linked to long-term respiratory outcomes, including recurrent wheeze and childhood-onset asthma^6–11^. Despite this substantial disease burden and long-term risk, there is limited understanding of the cell states and immune programs engaged in the pediatric airway during natural respiratory viral infection.

The nasal mucosa is a primary site of entry for most respiratory viruses and the location where host antiviral responses are first initiated^12,13^. At this interface, epithelial sensing, interferon (IFN) signaling, and immune cell recruitment shape early host-virus interactions that may influence subsequent immune responses and clinical outcomes. Studies of adult SARS-CoV-2 infection have shown that analysis of nasal mucosal responses reveals epithelial infection states and immune correlates of disease severity that are not evident from blood-based measurements alone, highlighting the nasal mucosa as a uniquely informative tissue for studying antiviral immunity at the site of infection^14–19^. However, the cellular organization of antiviral responses in the pediatric nasal mucosa during natural infection remains poorly characterized.

Although respiratory viral infections are ubiquitous in early childhood, different viruses are associated with distinct clinical phenotypes and disease trajectories in young children. Acute SARS-CoV-2 infection is typically mild during early childhood and is less likely to lead to severe respiratory illness or hospitalization; however, further studies are needed to understand its potential long-term, post-acute effects on children^20–23^. RSV is the most common cause of severe bronchiolitis, particularly in children younger than 6 months old^4,24^. Rhinovirus infection can also lead to bronchiolitis, but is most notable for its propensity to trigger childhood asthma exacerbations in susceptible hosts^9,11^. These divergent outcomes suggest that different viruses engage distinct host response programs at the mucosal interface. However, how epithelial and immune responses in the pediatric nasal mucosa differ across common respiratory viruses, and whether this heterogeneity relates to disease severity and symptoms such as wheezing, remains incompletely understood.

Progress in defining pediatric antiviral immunity has benefited from a range of complementary experimental approaches. *In vivo* single-cell studies of respiratory viral infection in children, largely focused on SARS-CoV-2 to date, have revealed unique features of pediatric immune responses, such as enhanced IFN responses and unique T cell populations^25–27^. However, most cohorts primarily include children older than 5 years or adolescents. *Ex vivo* air-liquid interface (ALI) cultures and organoid models have enabled further characterization of age-dependent antiviral responses^28–30^, identification of molecular features such as nucleolin and IGF1R as candidate receptors for RSV^31,32^, IL-1β-driven mucus production in the absence of IFN signaling in rhinovirus^33^, and investigation of viral effects on epithelial differentiation pathways^34–36^. While these systems provide important mechanistic insights under controlled conditions, they do not capture the full cellular heterogeneity or immune context present *in vivo*. In parallel, analyses of pediatric blood samples and tracheal aspirates from children with severe disease have revealed immune correlates of clinical outcomes, but provide limited information about epithelial infection states and early mucosal responses at the initial site of viral entry and do not capture the full spectrum of severity^37–41^. A recent upper-airway transcriptomic study of infants hospitalized with bronchiolitis of multiple viral etiologies, including RSV, identified compartmentalized mucosal immune signatures, including impaired interferon signaling and IL-36–associated neutrophilic inflammation, that correlate with disease severity but are not reflected in peripheral blood^42^. However, this study does not resolve epithelial infection states or define how epithelial and immune responses are coordinated within the nasal mucosa. Consequently, the *in vivo* identity of infected epithelial cells and the relationship between early nasal responses, disease severity, and wheezing outcomes remain incompletely defined.

Single-cell atlases provide a powerful framework for resolving cellular responses to infection *in vivo*, yet most existing atlases have been generated from cohorts in high-income settings^43–45^. This reflects the substantial logistical and technical challenges of generating high-resolution single-cell datasets in low– and middle-income countries, where the burden of severe pediatric respiratory infection is highest^3,5,46^. Whether cellular programs identified in single-cohort atlases are conserved across populations with distinct environments and disease burdens therefore remains largely untested. Addressing this gap will require the development of adaptable experimental and analytical approaches, as well as collaborative frameworks that support single-cell data generation and interpretation across diverse settings.

Here, we present a single-cell atlas of the pediatric nasal mucosa, profiling epithelial and immune cells from children under five years of age with SARS-CoV-2, rhinovirus, or RSV infection, alongside uninfected controls. By directly detecting viral transcripts within individual cells, we define virus-specific infected epithelial cell states, and through comparative analysis characterize distinct epithelial and immune response programs associated with each virus. We further relate nasal mucosal cell states to clinical outcomes, including hospitalization and wheezing illness, and contextualize genetic risk for childhood-onset asthma at the cellular level. Finally, we demonstrate the broader relevance of this atlas by applying it as a reference to independent pediatric cohorts from Kolkata, India, comprising healthy children, and rural Bangladesh, including healthy and infected children, revealing conserved cell states and virus-associated epithelial responses across geographically distinct populations. Together, this work provides a framework for understanding virus-specific antiviral immunity at the pediatric mucosal interface and establishes a broadly applicable reference for studying early-life respiratory disease.

## Results

### Developing an atlas of pediatric nasal mucosa antiviral responses

Respiratory viruses such as SARS-CoV-2, rhinovirus, and RSV are exceedingly common in young children and are known to cause disparate disease outcomes. We hypothesized that each virus might induce unique cellular responses in the pediatric nasal mucosa. To explore this, we collected research nasopharyngeal (NP) swabs from a convenience cohort of children visiting Boston hospitals (USA-Boston cohort) and performed single-cell RNA-sequencing (scRNA-seq) on 132 swabs from children younger than 5 years of age selected by viral positivity and representative of a range of clinical severity. 103 of the participants presented to the Boston Children’s Hospital (BCH) Emergency Department (ED) with acute respiratory symptoms. They were assigned to SARS-CoV-2, rhinovirus, RSV, or co-infection groups based on results from clinically indicated testing, laboratory-based RT-PCR of viral transport media from clinical samples, and/or RT-PCR of supernatants collected during processing of research NP swabs **(Figure 1A-B, Methods**). We also enrolled children younger than 5 years of age who presented to the BCH ED or an outpatient clinic at Massachusetts General Hospital (MGH) without respiratory symptoms as controls. When available, supernatants obtained during processing of research NP swabs were tested for influenza, SARS-CoV-2, rhinovirus, and RSV by RT-PCR. Controls with positive PCR values for any of these viruses and symptomatic patients with evidence of co-infections (PCR positive for more than one of SARS-CoV-2, rhinovirus, and RSV) were included in iterative clustering to define cell subsets, but were excluded from any virus-specific analysis. Our main cohort included 24 controls (BCH n=19, MGH n=5), 17 children with SARS-CoV-2, 39 children with rhinovirus, and 40 children with RSV **(Figure 1A, Supplementary Tables 1-2**).

**Figure 1:**
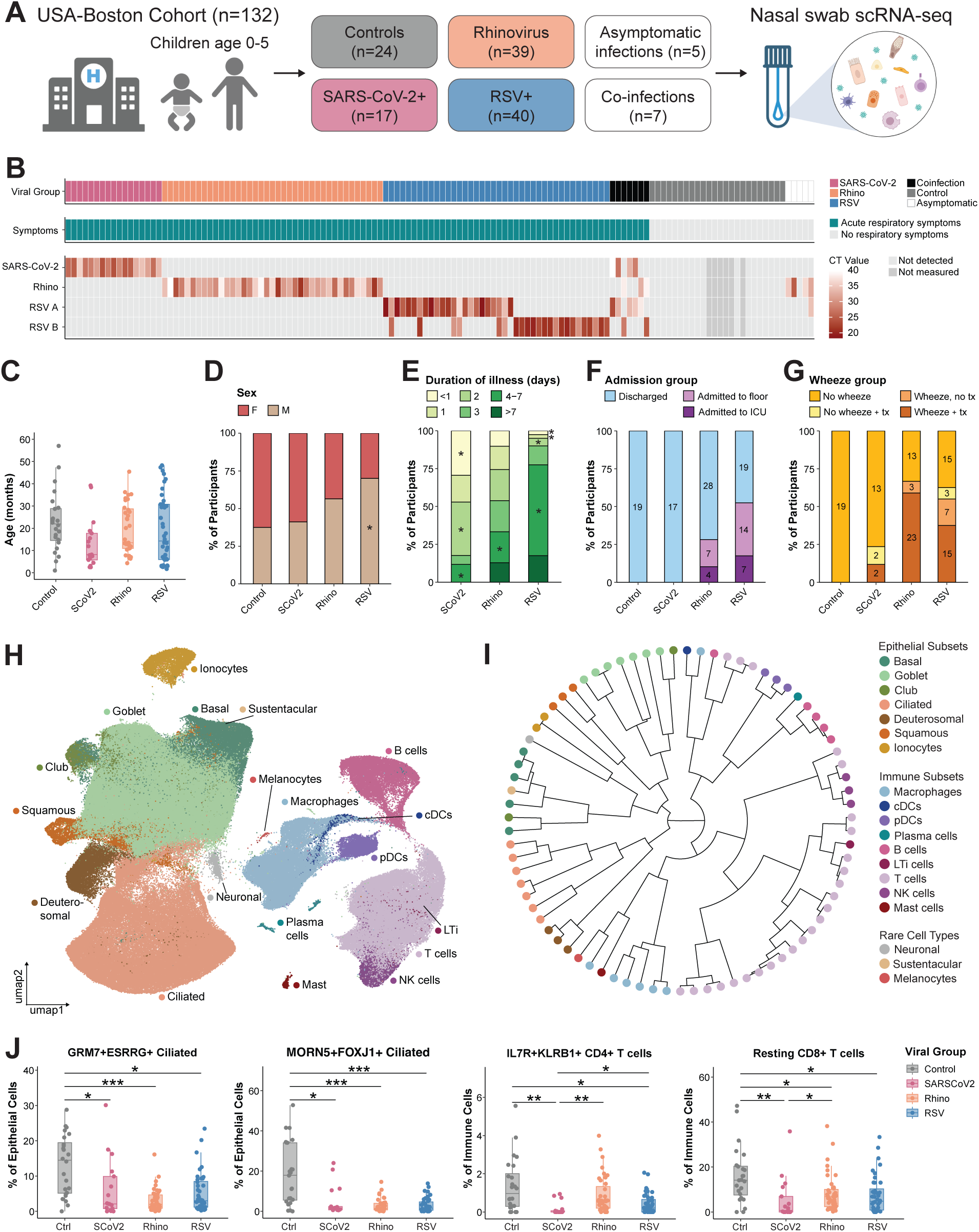
Cellular diversity in the pediatric nasal mucosa response to common respiratory viruses. **A.** Schematic of participant cohort and study design. RSV = respiratory syncytial virus; scRNA-seq = single-cell RNA sequencing. Participant metadata is summarized in **Supplementary Table 1** and provided in full in **Supplementary Table 2**. **B.** Heatmap of participant cohort colored by assigned viral group (top), the presence of acute respiratory symptoms (middle), and viral RNA cycle threshold (CT) value from supernatant collected during swab processing measured by qPCR (bottom). 1 asymptomatic participant was assigned based on a positive PCR from clinically sampled viral transport media. Not detected: CT value >40 or undetectable; Not measured: supernatant was not available; Rhino = rhinovirus. **C.** Age of each participant in months by viral group. SCoV2 = SARS-CoV-2. **D.** Distribution of sex within each viral group. *chi-sq. standardized Pearson residual > 2. **E.** Distribution of reported duration of illness within each viral group. *chi-sq. standardized Pearson residual > 2. **F.** Distribution of hospital admission within each viral group. Control samples are restricted to those collected at Boston Children’s Hospital (BCH). ICU = intensive care unit. **G.** Distribution of wheezing groups (see **Methods** for assignment criteria) within each viral group. Control samples are restricted to those collected at BCH. Tx = wheezing-directed medication. **H.** UMAP of all cells (n=335,174) from all participants colored by cell type. Cell type labels were assigned after iterative clustering. cDCs = conventional dendritic cells; pDCs = plasmacytoid dendritic cells; LTi = lymphoid tissue inducer cells; NK = natural killer. **I.** Hierarchical tree of all 79 cell subsets identified after iterative clustering. End nodes are colored by cell type of origin. Hierarchical distances were calculated from pseudobulk aggregates of each cell subset. See **Supplementary Figure 1G** for subset labels. **J.** Frequency of specified cell subsets as a percentage of either all epithelial or all immune cells per sample, compared across viral groups. Samples with <100 epithelial or immune cells were excluded. Ctrl = Control. **C,J**: Boxplots represent the median (center line), interquartile range (box), and 1.5x the interquartile range (whiskers). Statistical test is Kruskal-Wallis test with Benjamini-Hochberg correction for multiple comparisons. Dunn’s post-hoc test, *p<0.05, **p<0.001, ***p<0.0001.

There were no significant differences in age, sex, race, or ethnicity across viral groups, except for a higher proportion of males in the RSV group **(Figure 1C-D, Supplementary Table 1**). Two participants (1 SARS-CoV-2 case and 2 RSV) received RSV prophylaxis with the monoclonal antibody nirsevimab prior to their first RSV season and the mothers of 3 participants (1 SARS-CoV-2 and 2 RSV) had received RSV vaccination during pregnancy, though the majority of participants were enrolled prior to FDA approval of either preventative treatment **(Supplementary Table 1**). There were no significant differences in family history of atopy (asthma, eczema, or seasonal allergies), history of SARS-CoV-2 infection, or SARS-CoV-2 vaccination **(Supplementary Table 1**).

Acute SARS-CoV-2 infection generally causes mild symptoms in children^22^. In our cohort, children with SARS-CoV-2 had shorter duration of illness and less severe symptoms than those with rhinovirus or RSV **(Figure 1E, Supplementary Table 1**). RSV is the leading cause of bronchiolitis in young children and can cause both upper and lower respiratory disease^47^. In the group selected for scRNA-seq, children with RSV presented later in illness than children with SARS-CoV-2 or rhinovirus, had higher median modified BROSJOD bronchiolitis severity scores^48^ **(see Methods**), more frequent receipt of supplemental oxygen, and a higher rate of hospital admission **(Figure 1E-F, Supplementary Table 1**). Both rhinovirus and RSV are associated with persistent wheeze or asthma later in childhood^11^. Based on our interest in this pathology, we included children with wheezing on exam and/or treatment with systemic steroids, bronchodilators, or both in the groups selected for scRNA-seq **(Figure 1G, Supplementary Table 1**).

Viable epithelial and immune cells were isolated from each nasal swab as previously described by our group^15,19^. After iterative removal of low-quality cells, pool-specific clusters, and doublets, we obtained 335,174 high-quality cells. Because we did not observe notable differences in sample quality nor effects of technical or demographic factors on gene expression **(Supplementary Figure 1A-F, Methods**), and because we wanted to preserve biological variability across viral groups, we proceeded without data integration.

We performed iterative clustering to define the cell types and states represented in our dataset (Supplementary Figure 1G, Supplementary Figures 2-3, Supplementary Table 3). We identified diverse epithelial populations, including basal stem cells (*KRT5, KRT15*), intermediary club cells (*SCGB1A1, SCGB3A1*), mucus-producing goblet cells (*MUC5AC, BPIFA1*), and squamous cells (*SPRR3*, *SPRR2E*) important in barrier function **(Figure 1H, Supplementary Figure 2B**)^49–52^. There were also robust populations of ciliated epithelial cells (*FOXJ1*, *CAPS, DNAAF1*), deuterosomal cells (*CCNO*, *CDC20B*)^53,54^, and ionocytes (*CFTR, RARRES2*)^55,56^ **(Figure 1H, Supplementary Figure 2C**). Each major epithelial cell type contained multiple transcriptionally distinct subsets **(Figure 1I, Supplementary Figure 1G, Supplementary Figure 2, Supplementary Table 3**). For example, within goblet cells we identified a subset marked by *CST1*, a gene relevant in asthma pathobiology^57–60^. Squamous cells included a *KRT13+HOPX+* subset resembling hillock cells, an important cell subset in repair of injured airways^55,61^. Deuterosomal cells separated into populations that were more ciliated-like (*MORN2, PIFO, CROCC2*) versus those resembling goblet cells (*MUC5AC, SERPINB3*). Among ciliated epithelial cells, which are expected to be the main target of viral infections, we observed multiple interferon-stimulated (IFN-stim) subsets (*RSAD1, IFIT1, CXCL11*), an NF-kB responsive subset (*NFKBIA, TNFAIP3*), and a subset exhibiting mixed ciliated and squamous-like features (*KRT17, S100A9, LAMB3).* We observed two ciliated subsets defined by genes involved in cellular growth and cilia development (*GRM7+ ESRRG+* ciliated, *MORN5+FOXJ1+* ciliated) which were enriched in controls, indicating a consistent decline in healthy ciliated populations in all three infections **(Figure 1J**).

We also recovered diverse immune cell types, including macrophages (*TYROBP, FCER1G*), conventional type-1 dendritic cells (*CLEC9A*), plasmacytoid dendritic cells (pDCs) (*IRF7, GZMB*), B cells (*MS4A1, BANK1*), plasma cells (*MZB1, JCHAIN*), T cells (*CD3D, TRAC*), and NK cells (*GNLY, PRF1*) **(Figure 1H, Supplementary Figure 3**). Subclustering revealed additional rare subsets, including monocyte-derived dendritic cells (MoDC) (*GPR183, CD86*), mucosal-associated invariant T (MAIT) cells (*ZBTB16, KLRB1*, *SCL4A10*)^62,63^, and lymphoid tissue inducer (LTi) cells (*KIT, LTB*)^64^. T cells further separated into CD4+, CD8+, gamma-delta (γδ), and cycling subsets **(Supplementary Figure 3A, 3C, see Methods**). Across immune lineages, we observed both quiescent or ribosomal-high states (e.g., Ribo-hi B cells, resting CD4+ T cells) and more activated subsets (e.g., pro-inflammatory macrophages, *HAVCR2*+*CD38*+ CD4+ T cells, cytotoxic CD8+ T cells), highlighting the diversity of activation states captured in our samples **(Figure 1I, Supplementary Figure 1G, Supplementary Figure 3, Supplementary Table 3**). Control samples had higher frequencies of *IL7R*+*KLRB1*+ CD4+ T cells with a naive / memory phenotype and transcriptionally quiescent CD8+ T cells, suggesting that upon infection, the phenotype of nasal immune cells shifts to a more active inflammatory state **(Figure 1J**).

Additional rare populations, such as melanocytes (*PMEL*, *MLANA,* n=122, 0.036% of all cells), sustentacular cells (*ERMN, GPX6, CYP2A13*, n=113, 0.034% of all cells)^65^, and neuronal-like cells (*RBFOX1, CTNAP2, PTPRD*, n=1,830, 0.55% of all cells), were also detected **(Figure 1H-I**). In total, we defined 18 cell types and 79 cell subsets across the dataset. This represents the most comprehensive description of nasal cellular diversity in young children to date, enabling detailed exploration of the shared and unique responses to each respiratory virus.

### RSV and SARS-CoV-2 each induce highly specific infected epithelial cell states

Nasal epithelial cells are a primary target of respiratory viruses. Given the distinct transmission kinetics and symptoms associated with SARS-CoV-2, rhinovirus, and RSV, we hypothesized that the intrinsic transcriptional response of infected epithelial cells may differ for each virus. We first quantified viral RNA within each cell. Using our previously described method^15,19^, we aligned our transcriptional data to a custom reference including human, SARS-CoV-2^66^, numerous rhinovirus genomes across species A, B, and C **(see Methods**), RSV-A^67^, and RSV-B^68^ genomes and assigned cells as viral RNA+ if the viral counts were above the expected background amount **(Supplementary Figure 4A, “alignment method”**). For SARS-CoV-2 and RSV, we recovered significant amounts of viral RNA among epithelial cells **(Supplementary Figure 4B**). For rhinovirus, very few transcripts were detected **(Supplementary Figure 4B**), and only aligned to Rhinovirus C^69^. We detected 4,465 SARS-CoV-2 RNA+, 150 rhinovirus RNA+, and 8,767 RSV RNA+ cells across the dataset. For SARS-CoV-2 and RSV, viral UMI detected in the single cell data and the frequency of viral RNA+ cells for each participant correlated strongly to the qRT-PCR CT value for viral RNA detected in the swab supernatant **(Supplementary Figure 4C**). We did not observe a correlation between viral transcripts detected and severity score for any of the viruses **(Supplementary Figure 4C**). Transcripts were detected across the SARS-CoV-2 and RSV viral genomes, and for RSV, participants were consistently found to have reads either from RSV-A or RSV-B **(Supplementary Figure 4D**).

Because very few rhinovirus RNA+ cells were identified by alignment to custom reference genomes, we considered that detection by this method may be hampered by the extensive viral diversity within rhinovirus, which has three different species and more than 170 different serotypes. As such, we applied the Kraken2^70^ metagenomics method on a single cell basis to quantify microbial reads in an unbiased manner **(Supplementary Figure 4A, “Kraken method”**). There was a strong correlation between SARS-CoV-2 and RSV viral+ cells between the Kraken and alignment methods **(Supplementary Figure 4E**). While a few more participants had detectable rhinovirus+ cells using this method, the frequency of viral RNA+ cells for rhinovirus was much lower than RSV or SARS-CoV-2 by either method **(Supplementary Figure 4E**). This method also permitted detection of other common respiratory viruses, including parainfluenza (276 positive cells total) and human metapneumovirus (HMPV) (141 positive cells total), across multiple participant groups **(Supplementary Figure 4F**). However, these detections were sparse and HMPV detection was distributed across groups, and therefore we did not reclassify or exclude participants based on these findings. Future studies with greater numbers of parainfluenza and HMPV infected participants are warranted to permit detailed analyses of infected cells and responses.

Next, we mapped viral RNA+ cells to defined epithelial cell subsets to evaluate the specific phenotypes of presumably infected cells **(Figure 2A-B**). SARS-CoV-2 and RSV viral RNA+ cells occupied distinct areas of the epithelial UMAP **(Figure 2B**), and a discrete epithelial subset comprised the majority of viral RNA+ cells for both viruses. In SARS-CoV-2, the primary viral RNA+ subset was *NFKBIA*+*ATF3*+ ciliated cells **(Figure 2C**), which were also enriched in SARS-CoV-2 cases **(Figure 2D**). *KRT17*+*S100A9*+ ciliated cells were the primary infected cell subset in RSV **(Figure 2C**) and were specific to RSV cases **(Figure 2D**). A *CXCL11*+ IFN-stim ciliated subset included cells containing transcripts for either SARS-CoV-2 or RSV **(Figure 2C**), and was increased in frequency compared to controls for both viruses **(Figure 2D),** although only a minority of cells in this subset were viral RNA+ (**Figure 2C**). These results suggest that while some RSV and SARS-CoV-2 infected cells share a similar phenotype (*CXCL11+* IFN-stim) the majority of epithelial cells infected by each virus exhibit distinct and unique transcriptional phenotypes. Assessment of subsets comprising viral RNA+ cells identified via the Kraken method demonstrated similar results, with broader detection in SARS-CoV-2 **(Supplementary Figure 4G**) and detection of rhinovirus+ cells in the *CXCL11+* IFN-stim ciliated subset. We observed minimal detection of viral transcripts across immune subsets with either method, with the highest amount in macrophages in samples obtained from patients in the SARS-CoV-2 group **(Supplementary Figure 4H**). In keeping with this, SARS-CoV-2 and RSV viral UMIs were lower in immune than in epithelial cells **(Supplementary Figure 4B**). Interestingly, the distribution of viral RNA+ cells did not correlate with expression of known viral receptors or host processing genes for SARS-CoV-2 and RSV **(Supplementary Figure 4I**), which may indicate a later stage of infection.

**Figure 2:**
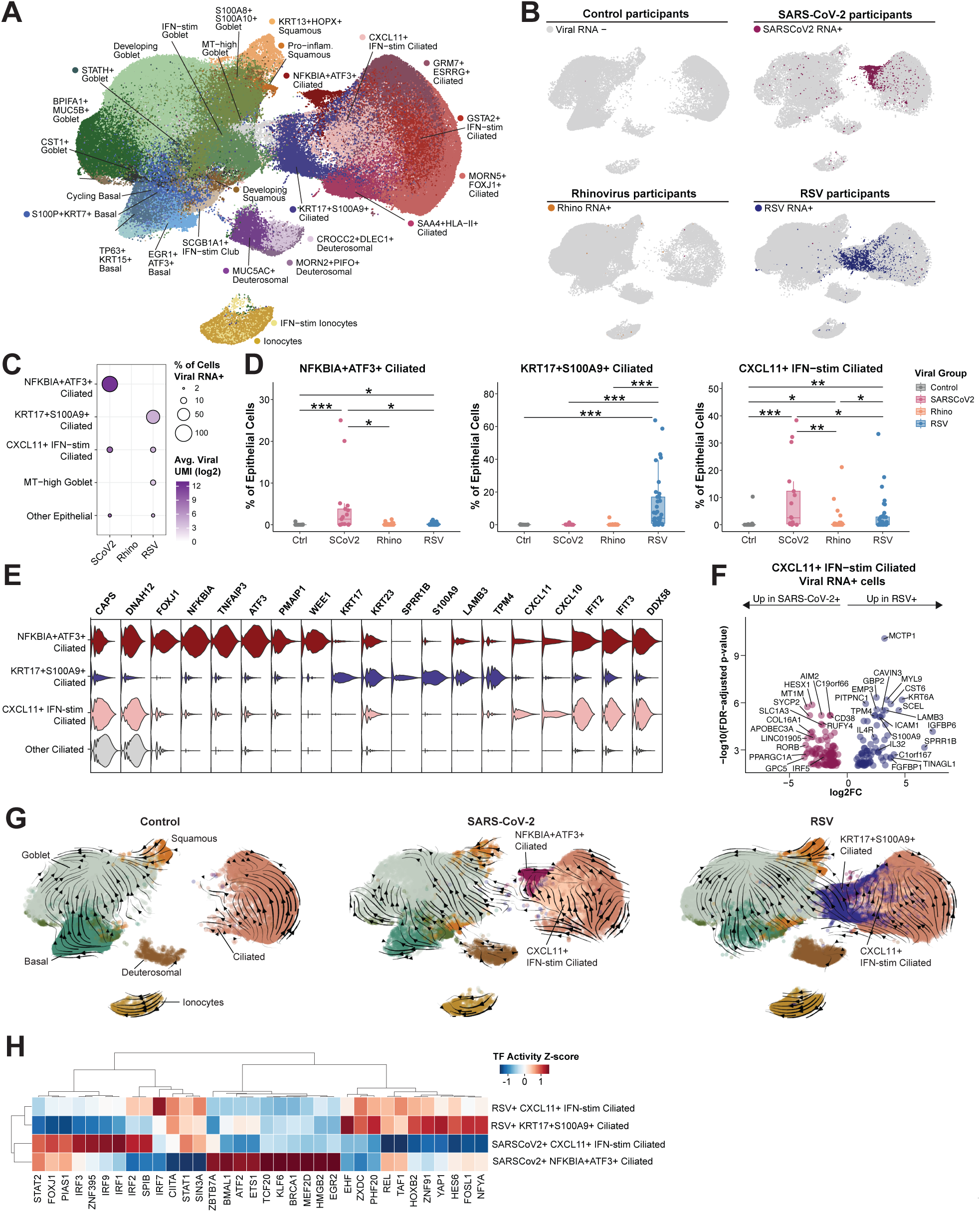
RSV and SARS-CoV-2 each induce highly specific infected epithelial cell states. **A.** UMAP of epithelial cells (n=224,431) from participants in control and monoinfection groups colored by cell subset. Pro-inflam = pro-inflammatory; MT-high = mitochondrial-high; IFN-stim = interferon-stimulated. **B.** Split UMAP of epithelial cells from each viral group (Control: n=34,031; SARS-CoV-2: n=39,836; Rhinovirus: n=61,198; RSV: n=89,366) colored by viral RNA status. **C.** Dotplot of viral RNA+ cells in each epithelial subset for SARS-CoV-2, rhinovirus, and RSV. Dot size represents the percentage of cells in the subset that are viral RNA+. Color represents the average viral transcripts (UMI) across all cells in the subset. Epithelial subsets with >5% of cells viral RNA+ were included, other subsets were collapsed. SCoV2 = SARS-CoV-2. **D.** Frequency of specified viral RNA+ cell subsets as a percentage of epithelial cells per sample, compared across viral groups. Samples with <100 epithelial cells were excluded. Boxplots represent the median (center line), interquartile range (box), and 1.5x the interquartile range (whiskers). Statistical test is Kruskal-Wallis test with Benjamini-Hochberg correction for multiple comparisons. Dunn’s post-hoc test, *p<0.05, **p<0.001, ***p<0.0001. Ctrl = Control; Rhino = Rhinovirus. **E.** Violin plot of marker genes for select viral RNA+ epithelial subsets. **F.** Differentially expressed (DE) genes between RSV RNA+ cells and SARS-CoV-2 RNA+ cells within the *CXCL11*+IFN-stim ciliated subset. Participants with at least 20 RNA+ *CXCL11*+IFN-stim ciliated cells (SARS-CoV-2: n=4; RSV: n=6) were included in pseudobulk DE analysis. Wald significance test with Benjamini-Hochberg correction. Genes with adjusted p<0.05 are shown. Full list of DE genes is provided in **Supplementary Table 4**. **G.** RNA velocity streams for all epithelial cells split by viral group for Control, SARS-CoV-2, and RSV groups, colored by cell type with *NFKBIA+ATF3+* ciliated, *CXCL11*+IFN-stim ciliated, and *KRT17+S100A9+* ciliated subsets highlighted. **H.** Heat map of inferred transcription factor (TF) activity, z-score normalized between groups of cells, for select viral RNA+ cells, split by *NFKBIA+ATF3+* ciliated, *CXCL11+*IFN-stim ciliated, and *KRT17+S100A9+* ciliated subsets, showing the top 10 transcription factors for each group of cells.

We further explored the phenotypes of the viral RNA+ cells. In addition to the negative regulators of NF-kB signaling *NFKBIA* and *ATF3*, the SARS-CoV-2 specific subset expressed many regulators of cell growth and apoptosis, including *ATF3, PMAIP1, and WEE1* **(Figure 2E**). These cells resembled a SARS-CoV-2 RNA-high subset defined by the circadian clock gene *PER1* and growth regulator *EGR1* that we previously identified as consistently enriched in viral RNA+ cells across SARS-CoV-2 variants^19^, as well as a “hyperinfected” ciliated cell subset described in a SARS-CoV-2 human challenge study^18^. Differentially expressed genes identified in these previously described subsets shared substantial overlap with those identified within *NFKBIA*+*ATF3*+ ciliated cells in this dataset, indicating a conserved signature of SARS-CoV-2 infected nasal epithelial cells **(Supplementary Figure 4J**).

The RSV-specific subset was defined by specific cytokeratins (*KRT17, KRT23*), as well as genes associated with squamous differentiation and adhesion (*SPRR1B, LAMB3, TPM4*) **(Figure 2E**). Since ciliated cells are thought to be the main target of RSV, the downregulation of ciliated markers and upregulation of squamous markers was unexpected. When we evaluated the differentially expressed genes between SARS-CoV-2 RNA+ and RSV RNA+ cells within the shared *CXCL11*+ IFN-stim subset, we saw that the RSV RNA+ cells upregulated genes from the same squamous-like program, including *LAMB3, SPRR1B, S100A9, and IGFBP6*, suggesting even within the same subset, RSV+ cells adopt a unique transcriptional phenotype **(Figure 2F, Supplementary Table 4**). Despite the differences in phenotypes between RSV and SARS-CoV-2 infected cells, RNA velocity analyses were consistent with the hypothesis that ciliated epithelial cells are the precursors to both viral-specific populations **(Figure 2G**).

Investigation of transcription factor (TF) activity demonstrated unique patterns in each viral RNA+ subset. The SARS-CoV-2 specific subset was enriched for transcription factors controlling cell differentiation and apoptosis (*ATF2, KLF6*) **(Figure 2H**). Activity of canonical IFN-responsive transcription factors (*IRF1, IRF3, IRF7, IRF9*) was highest in *CXCL11+* IFN-stim ciliated cells. Differential enrichment of STAT family members was observed, with higher inferred *STAT2* activity in *NFKBIA+ATF3*+ ciliated cells and higher *STAT1* activity in *KRT17+S100A9+* ciliated cells **(Figure 2H**). Given that *STAT1* and *STAT2* combined are required for IFNα or IFNλ signaling, while *STAT1* homodimers form downstream of IFNγ, this result could indicate stronger type-II IFN activation in RSV infected epithelial cells than in SARS-CoV-2 infected cells. The RSV-specific *KRT17*+*S100A9*+ subset was also uniquely enriched for transcription factors promoting cell growth and epithelial-mesenchymal transition (*EHF*, *YAP1, FOSL1*) **(Figure 2H**). Together, these analyses demonstrate that SARS-CoV-2 and RSV each induce a distinct infected ciliated cell state defined by virus-specific transcriptional regulatory programs.

### SARS-CoV-2 induces a more robust IFN response compared to rhinovirus and RSV

The detection of viruses through pattern recognition receptors and subsequent production of IFNs to warn neighboring cells are essential steps in containing viral infections. Our group and others have demonstrated that in adults, a robust upper airway IFN response is associated with milder COVID-19 symptoms^15,71^. Multiple groups have compared IFN responses to SARS-CoV-2 between children and adults, and suggested that one reason children tend to have milder COVID-19 symptoms is a stronger initial IFN response^25,26,28^. We therefore wondered whether a robust nasal IFN response was a universal feature of childhood respiratory infections, or if it varied across viruses. We first examined expression of genes encoding pattern recognition receptors (PRRs) and associated signaling proteins. We found that these genes were expressed in all three viral RNA+ subsets, though to a lower degree in *KRT17*+*S100A9*+ ciliated cells **(Supplementary Figure 5A**).

Next, we defined the main producers of IFNs in our dataset. Among epithelial cells, the viral RNA+ subsets were the only subsets in which we detected type I (*IFNB1, IFNE*) or type III (*IFNL1, IFNL2, IFNL3)* interferons **(Figure 3A**). When we evaluated the differentially expressed genes between RSV RNA+ and RSV RNA-cells in the RSV-specific *KRT17*+*S100A9*+ ciliated subset, we similarly found that type III IFN genes were upregulated among viral RNA+ cells **(Figure 3B, Supplementary Table 4**). This analysis was not possible in the *NFKBIA*+*ATF3*+ ciliated subset because there were not sufficient numbers of viral RNA-cells. Among immune cells, a subset of plasmacytoid dendritic cells (IFN-high pDCs) were the primary producers of type I and type III IFNs **(Figure 3C**). This subset, though rare, was highest in frequency in SARS-CoV-2 cases **(Figure 3D**) and SARS-CoV-2 participants for whom this subset was present had higher viral loads **(Supplementary Figure 5B**). We next evaluated the response to type I IFN signaling using a signature derived from IFNα cytokine treatment of nasal epithelial cells **(Supplementary Table 5**)^72,73^. Scoring pseudobulked epithelial cells from each participant with this signature revealed significantly higher IFNα response pathway activity in SARS-CoV-2 cases compared to both controls and other viral groups **(Figure 3E, Supplementary Figure 5C**). The epithelial IFNα response was significantly correlated with viral load only in RSV cases, and was not associated with age, duration of infection at time of sampling, or disease severity **(Figure 3F**).

**Figure 3:**
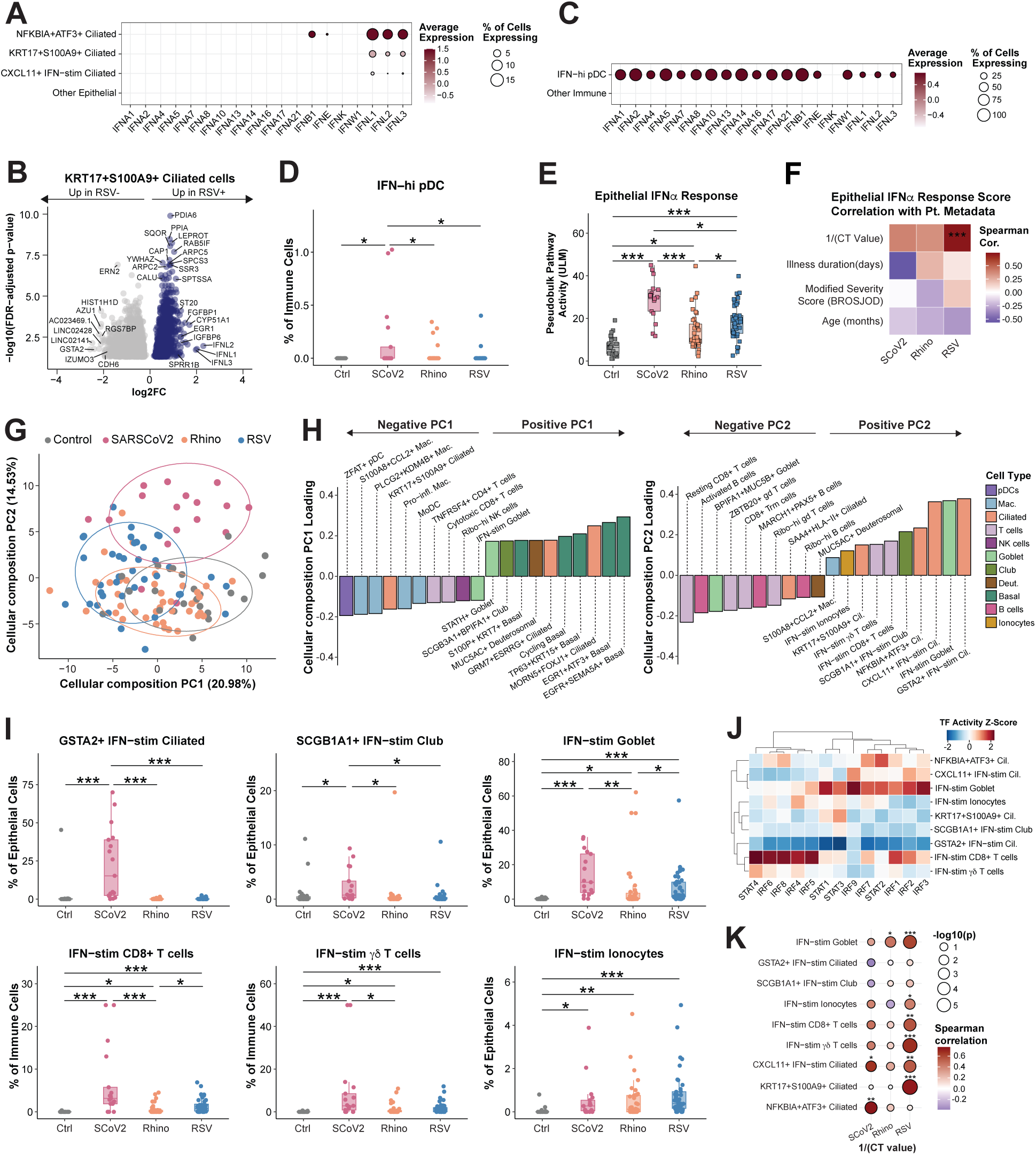
SARS-CoV-2 induces a more robust IFN response compared to rhinovirus and RSV. **A.** Dotplot of IFN gene expression by epithelial subset. Epithelial subsets with >1% of cells expressing any IFN gene were included, other subsets were collapsed. **B.** Differentially expressed (DE) genes between RSV RNA+ cells and RSV RNA-cells within the *KRT17+S100A9+* ciliated subset. Participants with at least 20 *KRT17+S100A9+* ciliated cells (n=26) were included in pseudobulk DE analysis. Wald significance test with Benjamini-Hochberg correction. Genes with adjusted p<0.05 are shown. Full list of DE genes is provided in **Supplementary Table 4**. **C.** Dotplot of IFN gene expression by immune subset. Immune subsets with >2% of cells expressing any IFN gene were included, other subsets were collapsed. **D.** Frequency of IFN-hi plasmacytoid dendritic cells (pDCs) as a percentage of immune cells per sample, compared across viral groups. Ctrl = Control; SCoV2 = SARS-CoV-2; Rhino = Rhinovirus. **E.** Pseudobulk pathway activity score per participant for interferon-alpha (IFNα) response in epithelial cells, compared across viral groups. ULM = univariate linear model. Signature provided in **Supplementary Table 5**. **F.** Spearman correlation between pseudobulk epithelial IFNα pathway activity score and select participant metadata variables within each viral group. CT value represents viral RNA PCR from scRNA-seq swab supernatants. **G.** Principal component analysis (PCA) of cellular composition for each sample. PCA was run on center-log-ratio (CLR) normalized cell subset abundances with percent of variance explained by each PC labeled on x and y axes. Ellipses for each viral group are centered at centroid and represent 80% confidence region. **H.** Cell subsets ranked by loadings for principal components 1 (left) and 2 (right). Top 10 positive and negatively ranked subsets are shown. Mac. = macrophage; Cil. = ciliated; Deut. = deuterosomal. **I.** Frequency of specified IFN-responsive epithelial and immune subsets as a percentage of immune or epithelial cells per sample, compared across viral groups. **J.** Transcription factor (TF) activity z-scores for select TFs across select IFN-responsive epithelial and immune subsets. **K.** Spearman correlation between CLR-normalized abundance of select cell subsets and inverse CT value from viral RNA PCR of scRNA-seq swab supernatants for each virus. **D,I:** Participants with <100 immune or epithelial cells were excluded. **D,E,I:** Boxplots represent the median (center line), interquartile range (box), and 1.5x the interquartile range (whiskers). Statistical test is Kruskal-Wallis test with Benjamini-Hochberg correction for multiple comparisons. Dunn’s post-hoc test, *p<0.05, **p<0.001, ***p<0.0001.

To assess the response to each virus in a more holistic manner, we turned to compositional methods that enable quantification of the cellular ecosystem of each sample^74,75^. Principal component analysis based on the cellular compositions revealed that rhinovirus cases were most similar to controls, RSV cases overlapped with both rhinovirus and controls but had some separation along PC1, and SARS-CoV-2 cases were the most distinct from other samples compositionally **(Figure 3G**). The subsets defining control cases (positive PC1, negative PC2) included basal cells, healthy ciliated cells, and resting immune populations as expected **(Figure 3H**). The main drivers of variation separating the RSV cases (negative PC1) were the *KRT17*+*S100A9*+ ciliated cells, along with multiple macrophage and T cell subsets, suggesting a heightened inflammatory response **(Figure 3H**). The cellular composition of SARS-CoV-2 cases (positive PC2) were defined by multiple IFN-responsive subsets **(Figure 3H**), including the *CXCL11*+ IFN-stim and *NFKBIA*+*ATF3*+ virally infected ciliated subsets. When we compared the frequency of IFN-responsive subsets identified through the compositional analysis, we observed that these subsets were either specific to SARS-CoV-2 (*GSTA2*+IFN-stim ciliated and *SCGB1A1*+IFN-stim club), increased in frequency in all viruses but to the greatest extent in SARS-CoV-2 (IFN-stim goblet, IFN-stim CD8+ T, IFN-stim γδ T), or in one case non-specifically enriched in all three viruses (IFN-stim ionocytes) **(Figure 3I**).

Noting that some IFN-responsive subsets were virus-specific while others were shared between viruses, we hypothesized that different transcription factors were activated across subsets. We next evaluated the use of key antiviral transcription factors across IFN-stim cell subsets. Transcription factors downstream of viral pattern recognition and IFN signaling, including *IRF3, IRF7, IRF1, IRF2,* and the ISGF3 complex (*STAT1/STAT2/IRF9*), showed the highest inferred activity in the same cell subsets, particularly IFN-stim goblet cells, IFN-stim CD8+ T cells, *CXCL11*+ IFN-stim ciliated cells, and the SARS-CoV-2-high *NFKBIA*+*ATF3*+ ciliated cells **(Figure 3J**)^76^. In contrast, the RSV-high subset *KRT17+S100A9+* ciliated cells and SARS-CoV-2-specific bystander population *GSTA2*+ ciliated cells had less activity of most IFN-response regulators except for *STAT1*. IFN-stim CD8+ T cells showed specific enrichment of *STAT4*, *IRF5*, *IRF6*, and *IRF8* activity, transcription factors previously implicated in IFN signaling, cytotoxic effector differentiation, and antiviral T-cell responses **(Figure 3J**)^76,77^. *STAT3* activity, which upregulates proinflammatory cytokine responses, was strongest in IFN-stim goblet and the RSV-high *KRT17*+*S100A9*+ ciliated cell subsets **(Figure 3J**).

We observed positive correlations between the abundance of multiple IFN-stim cell subsets and viral RNA levels in the sample, although the pattern differed by virus. In SARS-CoV-2, the viral RNA+ *NFKBIA+ATF3+* and *CXCL11*+ IFN-stim ciliated subsets correlated with viral load, whereas in rhinovirus only IFN-stim goblet cells showed such an association **(Figure 3K**). RSV demonstrated the greatest number of significant correlations between IFN-stim subsets and viral load **(Figure 3K),** further indicating that IFN activation in RSV is closely linked to the amount of virus present. Taken together, our data indicate that SARS-CoV-2 induces a significantly more robust productive IFN bystander response than either rhinovirus or RSV, which may reflect a more amplified antiviral state that is not solely proportional to viral burden. These results suggest that young children have a graded IFN response dependent on the specific viral challenge.

### Rhinovirus infection exhibits secretory-cell polarization of the epithelium

Given the diminished IFN responses in rhinovirus in comparison to RSV and particularly in comparison to SARS-CoV-2, we sought to further understand the epithelial response specific to rhinovirus. We shifted from subset-level frequencies to analysis at the broader epithelial cell-type level for each virus compared to controls, using centered log-ratio (CLR)-normalized abundance to assess compositional changes^74^. We found that in all three viruses, there was an increase in abundance in goblet cells **(Figure 4A**). In rhinovirus cases, there was a notable depletion of ciliated epithelial cells and an increase in deuterosomal cells **(Figure 4A**). Because ciliated cells are the primary targets of rhinovirus infection, we hypothesized that receptor-expressing cells might be selectively lost. While *ICAM1* and *LDLR*, the receptors used by rhinovirus-A and rhinovirus-B major and minor groups, respectively, were broadly expressed at low levels, CDHR3, the receptor for rhinovirus-C, was most highly expressed in ciliated cells **(Figure 4B**). Consistent with this distribution, rhinovirus cases exhibited a significant reduction in *CDHR3*+ epithelial cells compared to controls and other viral infections **(Figure 4C**). These findings suggest that rhinovirus-infected CDHR3+ ciliated cells may be lost through cell death or are underrepresented in our dataset, potentially explaining the limited detection of rhinovirus RNA+ cells.

**Figure 4:**
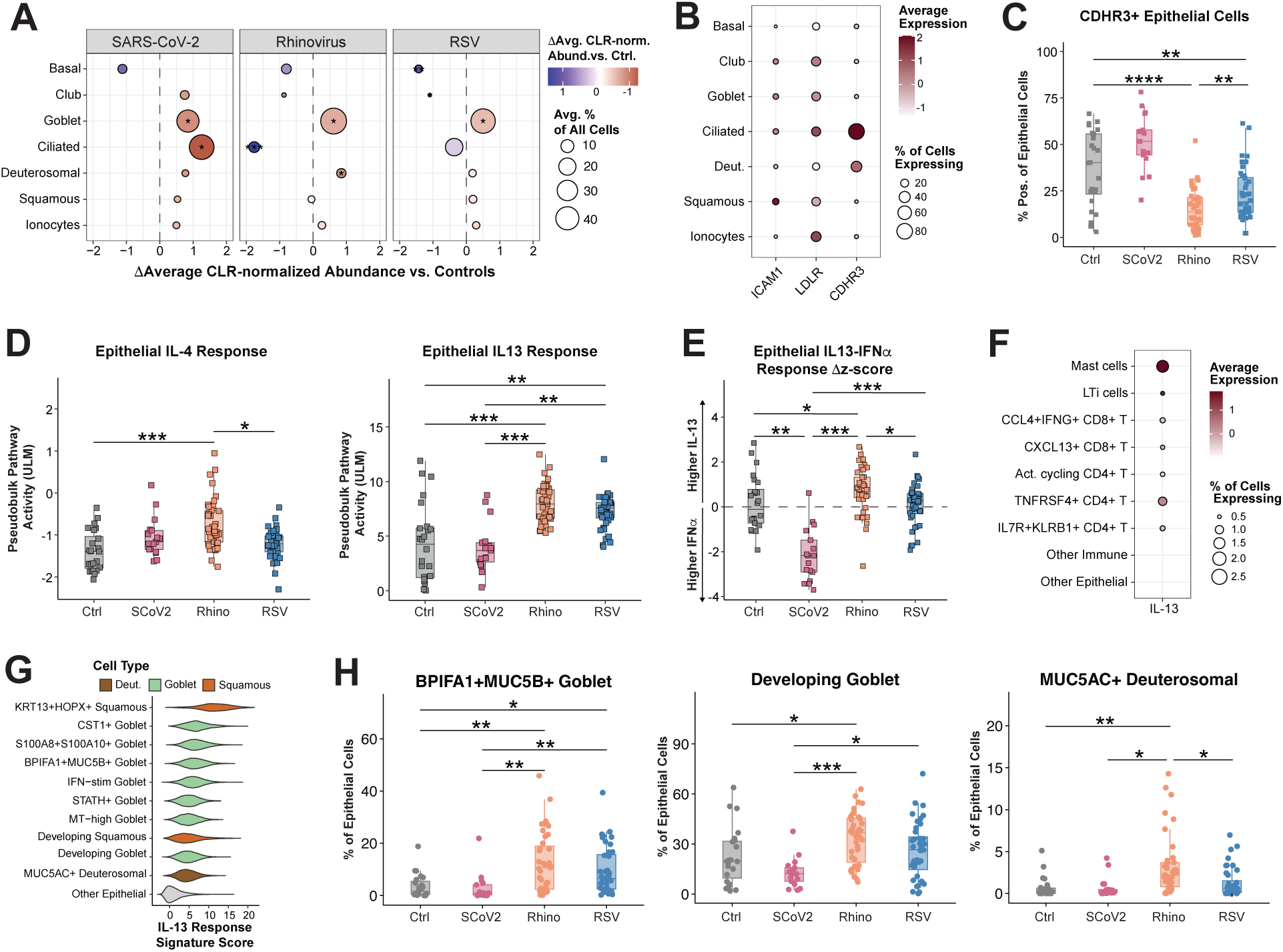
Rhinovirus is associated with secretory-cell polarization of the epithelium. **A.** Difference in average (Avg.) center-log-ratio (CLR)-normalized cell type abundance (Abund.) compared to controls (Ctrl.) for epithelial cell types. Dot size represents frequency of the cell type out of all cells in the sample, averaged per viral group. Statistical test is Wilcoxon test with Benjamin-Hochberg correction for multiple comparisons. *p<0.05, **p<0.01, ***p<0.001. **B.** Dotplot of rhinovirus receptor gene expression by epithelial cell type. **C.** Percentage of epithelial cells expressing the rhinovirus receptor gene *CDHR3* per participant, compared across viral groups. Cells were defined as CDHR3+ if they contained >1 *CHDR3* UMI. Statistical test is Wilcoxon test with Benjamin-Hochberg correction for multiple comparisons. *p<0.05, **p<0.01, ***p<0.001. Ctrl = Control; SCoV2 = SARS-CoV-2; Rhino = Rhinovirus. **D.** Pseudobulk pathway activity score per participant for IL-4 (left) and IL-13 (right) response genes in epithelial cells, compared across viral groups. ULM = univariate linear model. Signatures provided in **Supplementary Table 5**. **E.** Difference in scaled Z-scores of pseudobulk IL-13 and IFNα activity scores in epithelial cells per participant, compared across viral groups. **F.** Dotplot of IL-13 gene expression by cell subset. Subsets with >0.5% of cells expressing IL-13 or average scaled expression >2 were included, other subsets were collapsed. Act. = activated. **G.** Top 10 epithelial cell subsets by IL-13 response signature. Scoring was done on a downsampled object containing 10% of cells from each epithelial cell subset. **H.** Frequency of specified epithelial subsets as a percentage of epithelial cells per sample, compared across viral groups. Participants with <100 epithelial cells were excluded. **C,D,E,H:** Boxplots represent the median (center line), interquartile range (box), and 1.5x the interquartile range (whiskers). **D,E,H:** Statistical test is Kruskal-Wallis test with Benjamini-Hochberg correction for multiple comparisons. Dunn’s post-hoc test, *p<0.05, **p<0.001, ***p<0.0001.

Type 2 cytokines IL-4 and IL-13 are known to increase secretory and goblet cell differentiation. Furthermore, rhinovirus induces a strong type 2 response in some individuals^78–83^. When we evaluated response signatures for these cytokines **(Supplementary Table 5**)^72,73^, we observed a stronger IL-13 response in comparison to the IL-4 response **(Supplementary Figure 5C**). Comparison across viral groups revealed a modest but significant increase in IL-4 response scores in rhinovirus cases compared to RSV cases and controls **(Figure 4D**). IL-13 response scores were notably higher in both rhinovirus and RSV cases compared to controls and SARS-CoV-2 cases **(Figure 4D**). To evaluate the balance between type 1 and type 2 immune responses (represented by IFNα and IL-13, respectively) in each participant, we calculated the difference in the z-scores for each response signature. We observed a gradient in this ratio, with a strong enrichment in IFNα response in SARS-CoV-2, as noted in Figure 3, an even ratio for RSV, and an enrichment in IL-13 responses for rhinovirus **(Figure 4E**). Epithelial IL-13 response was not significantly associated with viral load, age, duration of illness, or severity **(Supplementary Figure 5D**).

We next evaluated which subsets were the main producers of IL-13. Although detection in general was quite low, we found that mast cells had the highest expression, followed by *TNFRSF4*+ CD4+ T cells **(Figure 4F**). Ranking of epithelial cell subsets by their IL-13 response score showed that *KRT13+HOPX+* squamous cells, the subset we noted resembles hillock cells, had the strongest score **(Figure 4G, Supplementary Figure 5C**). These were followed by multiple goblet subsets and *MUC5AC+* deuterosomal cells, indicating that IL-13 is likely acting on secretory cells in the epithelium. As deuterosomal cells are thought to be a differentiation intermediate towards the ciliated cell^53,54^, the expression of *MUC5AC* was unexpected and could indicate goblet cell polarization within this population. When we compared the frequency of the subsets with the highest IL-13 scores, we found that many were enriched in rhinovirus cases, including *BPIFA1+MUC5B+* goblet cells, developing goblet cells, *MUC5AC+* deuterosomal cells, and *CST1+* goblet cells **(Figure 4H, Supplementary Figure 5E).** Together, these results indicate a preferential secretory polarization during the immune response to rhinovirus infection.

### RSV is characterized by a heightened inflammatory and cytotoxic response

Immune cell infiltration is a common feature following respiratory viral infection of the nasal epithelium. However, particularly in RSV infection, the balance between protective and pathogenic inflammation may be determined by the quantity and phenotype of immune cells recruited to the airway^24^. Severe RSV infection has been linked to elevated numbers of neutrophils, monocytes, NK cells, T cells, and inflammatory cytokines such as IL-6 and IL-8 in the airways, as well as lower levels of type I and type II IFNs in blood and nasal secretions^40,84–88^. As such, we hypothesized that the nasal cellular immune response to RSV may be distinct from that of other viruses. We first assessed overall changes in CLR-normalized immune cell type abundance, and observed that in both rhinovirus and RSV cases, plasmacytoid dendritic cells (pDCs), NK cells, and cycling T cells were significantly increased **(Figure 5A**). In contrast, there were no significant increases in immune cell abundance in SARS-CoV-2 cases (**Figure 5A**). Next, we explored which cytokines, chemokines, and other immune signaling ligands **(see Methods, Supplementary Table 6**) were significantly upregulated compared to controls for each virus. We found that macrophages and CD8+ T cells were the strongest sources, and that despite limited changes in immune cell abundance, SARS-CoV-2 cases still had significant upregulation of many immune ligands **(Figure 5B, Supplementary Table 6**).

**Figure 5:**
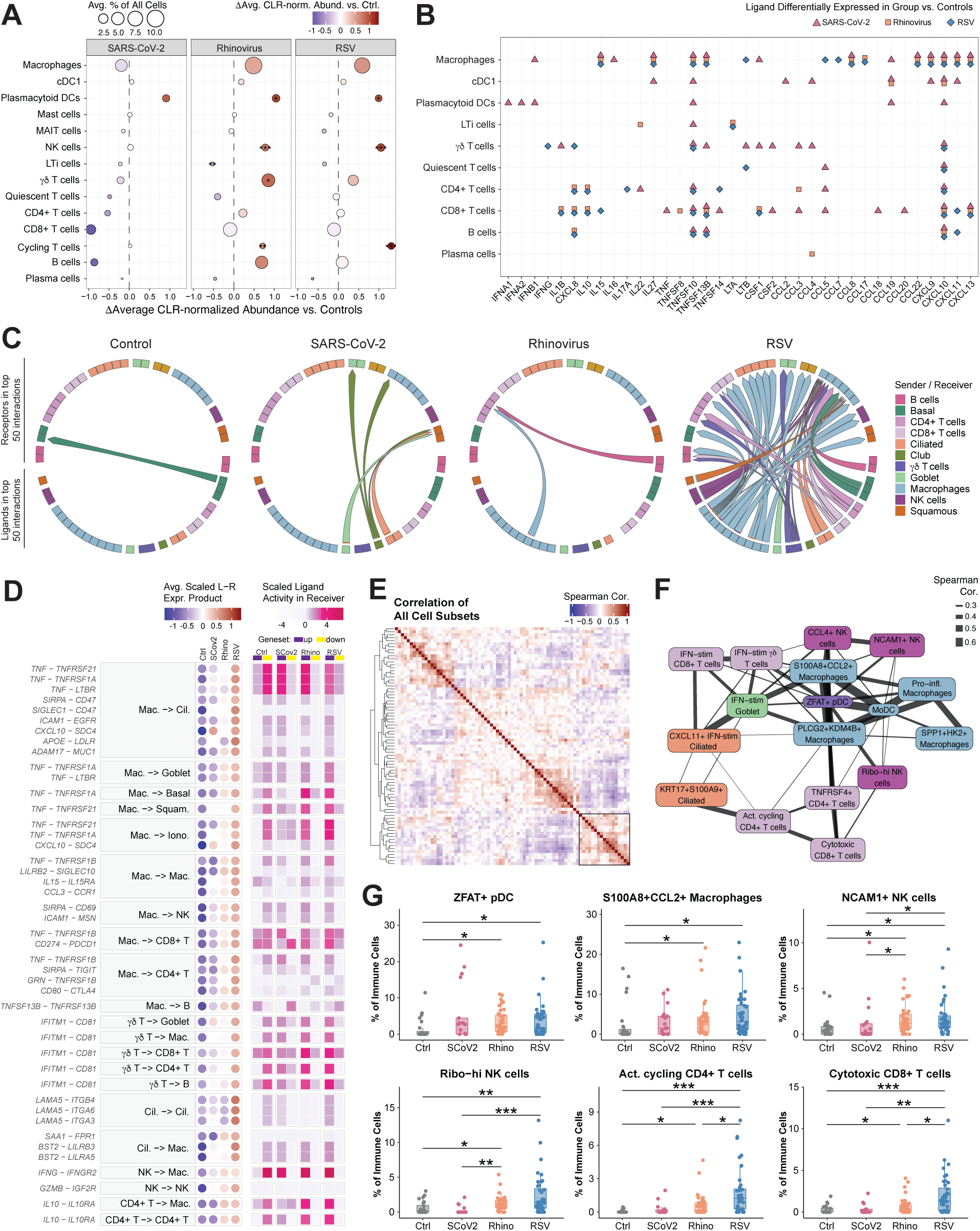
Nasal immune cell responses to viral infection reveal a heightened inflammatory and cytotoxic response in RSV. **A.** Difference in center-log-ratio (CLR)-normalized cell type abundance compared to controls for immune cell types. Dot size represents frequency of the cell type out of all cells in the sample, averaged per viral group. cDC1 = conventional type 1 dendritic cell; MAIT = mucosal-associated invariant T; NK = natural killer; LTi = lymphoid tissue inducer. **B.** Immune-signaling ligands per immune cell type differentially expressed (DE) relative to controls for each viral group. Cell types where at least 2 samples in every group had at least 5 cells were included. Cell type – ligand combinations with significant results (FDR <0.05) for at least one viral group are shown. See **Supplementary Table 6** for list of ligands evaluated. **C.** Chord diagram of top 50 overall ranked cell-cell interactions (multinichenet), divided by group specificity. Starting point of chord represents ligand and endpoint represents receptor. Chords are colored by sender cell type. See **Supplementary Figure 6A** for ligand and receptor labels and **Supplementary Table 6** for list of top 500 ranked interactions. **D.** Prioritization criteria for select cell-cell interactions in the RSV group within the top 100 overall. Interactions with sender cell types in the top 5% percent of interactions are shown. In cases where ligand-receptor-receiver combinations are not unique, interaction with the highest prioritization score is shown. Circles: scaled product of pseudobulk expression of ligand in sender and receptor in receiver, averaged across samples in the group. Squares: scaled ligand activity score in receiver cell type, representing enrichment of ligand target genes in differentially expressed (DE) genes; up: upregulated genes in that group, down: downregulated genes in that group. Ctrl = Control; SCoV2 = SARS-CoV-2; Rhino = Rhinovirus, Avg. Scaled L-R Expr. Product = average scaled ligand-receptor expression product. **E.** Spearman correlation (cor.) of CLR-normalized cell subset abundances among all samples. Both rows and columns were clustered using hierarchical clustering with Euclidean distance and complete linkage. **F.** Network plot of cell subsets highlighted in the box in **E**. Edge weight represents spearman correlation. Nodes are colored by cell type of origin. **G.** Frequency of specified immune subsets as a percentage of immune cells per sample, compared across viral groups. Participants with <100 immune cells were excluded. Boxplots represent the median (center line), interquartile range (box), and 1.5x the interquartile range (whiskers). Statistical test is Kruskal-Wallis test with Benjamini-Hochberg correction for multiple comparisons. Dunn’s post-hoc test, *p<0.05, **p<0.001, ***p<0.0001.

We next sought to broadly understand differences in cell-cell communication between groups. We applied multinichenet to rank the ligand-receptor interactions by 1) differential expression in each group, 2) consistency of expression across samples, 3) ligand activity enrichment in the receiver cell types, and 4) cell type specificity^89^. We found that of the top 50 ranked interactions, the majority were in the RSV group **(Figure 5C, Supplementary Figure 6A, Supplementary Table 6**) and that this was consistent across cutoffs **(Supplementary Figure 6B**). SARS-CoV-2 cases had the second-highest number of significant interactions and consistently strong prioritization scores **(Supplementary Figure 6C**). To further characterize the RSV-specific interactions, we selected those involving the top sender cell types, which included macrophages, γδ T, NK, ciliated, CD4+ T, and CD8+ T cells **(Supplementary Figure 6D**). Notably, macrophages emerged as a central communication hub, participating in a large fraction of the top-ranked interactions. The dominant signals included cytokines (*TNF, IL15, IL10, TNFSF13B*), IFN and IFN-related genes (*IFNG, CXCL10, IFITM1-CD81* interactions), and immune regulatory or adhesion-associated molecules (*SIRPA, ICAM1, SIGLEC1*). Although ligand activity scores were often comparable across groups, RSV cases consistently showed the highest expression of the corresponding ligand–receptor pairs **(Figure 5D**). Together, these findings indicate that RSV infection is marked by enhanced macrophage-centered inflammatory and interferon signaling, with greater immune-immune and immune-epithelial communication than SARS-CoV-2 or rhinovirus.

We next explored which cell subsets change in concert with each other. Correlation of the CLR-normalized abundances of all cell subsets^75^ revealed several clusters representing groups of coordinated cell subsets **(Figure 5E, Supplementary Figure 6E**). A cluster of interest contained both RSV-infected ciliated cell subsets (*KRT17+S100A9+* and *CXCL11*+IFN-stim) as well as multiple inflammatory macrophage, NK cell, and T cell subsets **(Figure 5F, Supplementary Figure 6E**), demonstrating that these immune cell subsets increased in relative abundance together with RSV RNA+ epithelial cells. Many of these subsets were also upregulated in RSV cases, including *ZFAT*+ pDCs, *S100A8+CCL2+* macrophages, *NCAM1+* NK cells, ribosomal-high NK cells, activated cycling CD4+ T cells, cytotoxic CD8+ T cells, *CCL4+* NK cells, *PLCG2+KDM4B+* macrophages, *SPP1+HK2+* macrophages, and *HAVCR2+CD38+* CD4*+* T cells **(Figure 5G, Supplementary Figure 6F**). These results suggest that RSV is associated with a distinct network of activated, cytotoxic, and inflammatory immune cells that is not observed in SARS-CoV-2 or rhinovirus.

### Distinct cellular programs in severe RSV infection and rhinovirus-induced wheeze

RSV and rhinovirus cause a broad spectrum of disease phenotypes in young children, while SARS-CoV-2 is most commonly associated with a mild upper respiratory infection or febrile illness in this population. RSV can cause severe lower respiratory tract infection in otherwise healthy young children, while rhinovirus is strongly associated with viral-induced wheeze and asthma exacerbations in genetically-susceptible individuals. However, it is not clear whether the nasal responses to either virus correlate with disease severity or wheezing response. We identified several significant correlations between nasal cell subset abundance and specific clinical variables **(Figure 6A**). Of note, when participants were split by viral group, most associations did not retain significance **(Supplementary Figure 7**). Resting and Trm-like CD8+ T cells positively correlated with age, consistent with our previous findings^90^. In keeping with the known association between IL-13 responses and asthma, two goblet cell populations that we identified as being IL-13 responsive (*BPIFA1+MUC5B+* and *CST1+*) were associated with wheezing symptoms and treatments **(Figure 6A**). Mast cells and a *CXCL13*+ CD8+ T cell subset also correlated with wheezing symptoms and treatments **(Figure 6A**). Multiple RSV-associated subsets were associated with disease severity and respiratory support, including *KRT17+S100A9+* ciliated cells, activated cycling CD4+ T cells, cytotoxic CD8+ T cells, and ribosomal-high NK cells **(Figure 6A**), in keeping with the RSV participants having generally more severe illness.

**Figure 6:**
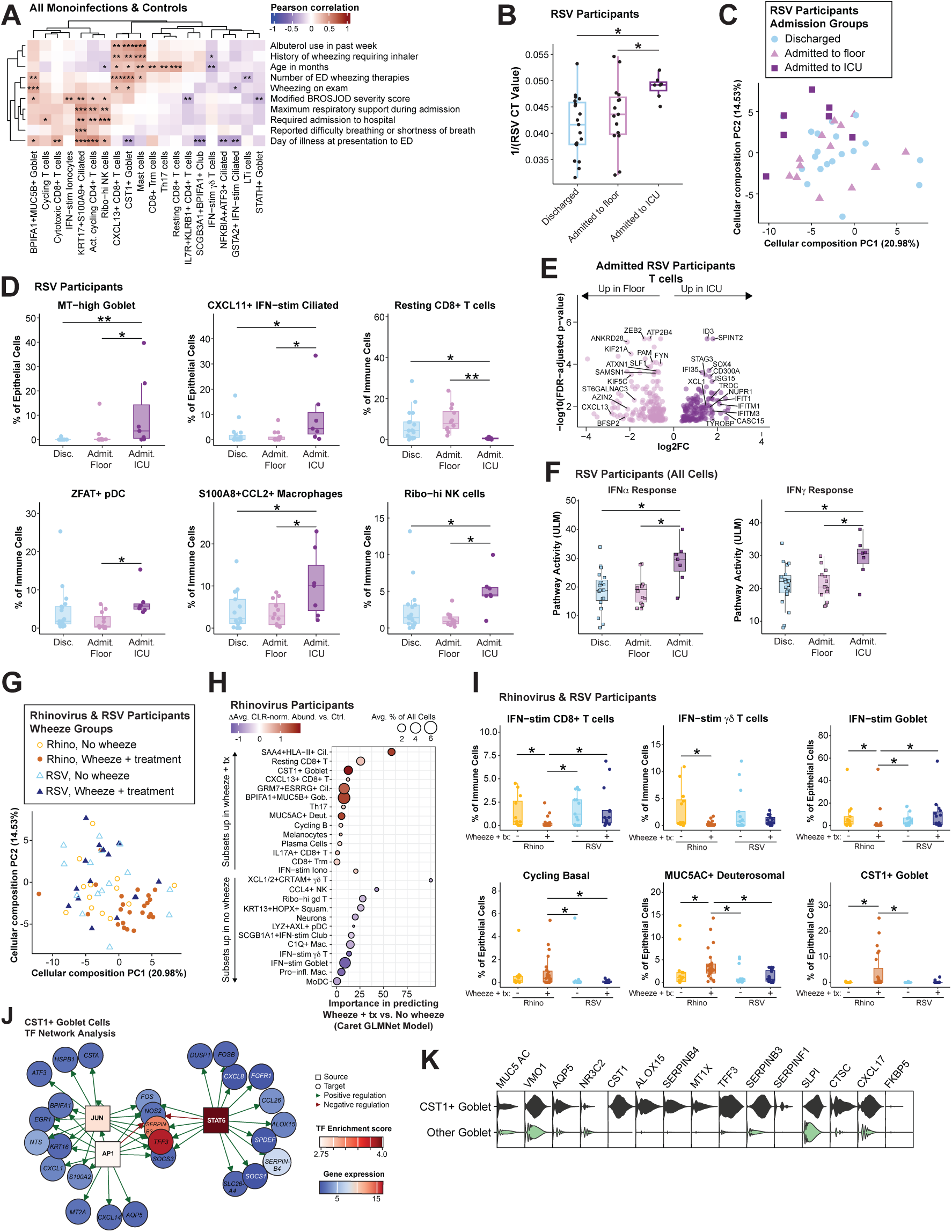
Cell states associated with severe RSV infection and rhinovirus-induced wheeze. **A.** Pearson correlation of CLR-normalized cell subset abundances among control, SARS-CoV-2, RSV, and rhinovirus participants. Both rows and columns were clustered using hierarchical clustering with Euclidean distance and complete linkage. P values from correlation test adjusted with Benjamini-Hochberg correction for multiple comparisons. *p<0.05, **p<0.01, ***p<0.001. ED = emergency department. **B.** Viral RNA level (1/CT value) from PCR of scRNA-seq swab supernatants for each sample in the RSV group divided by hospital admission status. ICU = intensive care unit. **C.** Principal component analysis (PCA) of cellular composition for each sample in RSV participants colored by admission groups. The percent of variance explained by each PC is labeled on x and y axes. PCA was run on center-log-ratio (CLR) normalized cell subset abundances. See **Figure 3G** for PC loadings. **D.** Frequency of specified subsets as a percentage of epithelial or immune cells in RSV participants compared across admission groups. Disc. = Discharged, Admit. floor = Admitted to floor, Admit ICU = Admitted to ICU. **E.** Differentially expressed (DE) genes in all T cells between RSV participants admitted to ICU or admitted to the floor by pseudobulk differential expression. Wald significance test with Benjamini-Hochberg correction. Genes with adjusted p value of <0.05 are displayed. Full list of DE genes is provided in **Supplementary Table 4**. **F.** Pseudobulk pathway activity score for IFNɑ (left) and IFNɣ (right) response genes per RSV participant in all cells, compared across admission groups. ULM = univariate linear model. Signatures provided in **Supplementary Table 5**. **G.** PCA of cellular composition for RSV and Rhinovirus participants colored by wheezing group (see **Figure 1E** for group breakdown). Treatment included nebulized bronchodilator and/or corticosteroids. **H.** Machine-learning-derived subset importance for distinguishing no wheeze vs wheeze + treatment for Rhinovirus participants by Caret’s penalized generalized linear (GLMNet) model. Color of dot indicates the difference in center-log-ratio (CLR)-normalized cell subset abundance between patients with wheeze+tx compared to no wheeze. Dot size represents frequency of the cell subset, averaged across all included rhinovirus participants. All subsets that contributed to model importance are shown. Cil. = ciliated; Gob. = goblet; Deut. = deuterosomal; Iono. = ionocytes; Squam = squamous; Mac. = macrophages. **I.** Frequency of specified subsets as a percentage of epithelial or immune cells for RSV and Rhinovirus participants categorized as having no wheeze/no treatment or wheeze + treatment. Tx = treatment; Rhino = rhinovirus. **J.** Transcription factor (TF) network analysis for the top 3 transcription factors within the *CST1*+ goblet cell subset. Boxes: transcription factors colored by enrichment score, circles: target genes colored by gene expression relative to all epithelial cells. **K.** Violin plot of marker genes for *CST1+* goblet compared to all other goblet cells. **D,I:** Participants with <100 epithelial or immune cells were excluded. **B,D,F,I:** Boxplots represent the median (center line), interquartile range (box), and 1.5x the interquartile range (whiskers). Statistical test is Kruskal-Wallis test with Benjamini-Hochberg correction for multiple comparisons. Dunn’s post-hoc test, *p<0.05, **p<0.001, ***p<0.0001

We next compared the nasal mucosa responses to RSV between participants who were discharged from the ED, admitted to the hospital for general care on the floor, or admitted to the ICU for intensive respiratory support. ICU cases had higher overall viral loads in the nasal swab sample **(Figure 6B**) and trends towards higher amounts of viral RNA detected at the single-cell level and frequency of viral RNA+ cells **(Supplementary Figure 8A-B**). We next examined the cellular composition PCA (as in **Figure 3F-G**) by the RSV admission groups. Participants who were admitted to the ICU separated along PC2, suggesting increased abundance of IFN-responsive subsets **(Figure 6C**, **Figure 3G**). RSV cases who were admitted to the ICU were characterized by increased frequencies of MT-high goblet cells, *CXCL11*+ IFN-stim ciliated cells, *ZFAT*+pDCs, *S100A8+* macrophages, and ribosomal-hi NK cells, as well as a decline in resting CD8+ T cells **(Figure 6D**). As the MT-high goblet cells and *CXCL11+* IFN-stim ciliated cells both contain some RSV RNA+ cells **(Figure 2C**), this result suggests greater RSV dissemination amongst the epithelium in more severe disease. Further investigation of differentially expressed genes in T cells between RSV cases admitted to the floor and intensive care demonstrated upregulation of many IFN-stimulated genes (*IFIT1, IFITM1, IFITM3, ISG15, IFI35*) in ICU cases **(Figure 6E**). Gene activity signatures **(Supplementary Table 5**) of IFNα and IFNγ were also significantly higher in ICU cases compared to either discharged or admitted to floor participants **(Figure 6F**). We did not observe a particular pattern in cellular composition nor any changes in IFN response scores in rhinovirus cases divided by admission status **(Supplementary Figure 8C-D**). Together, these results indicate that severe RSV disease is characterized by a high nasal viral load at the time of ED presentation, a strong type I / type II IFN response, and increases in inflammatory immune subsets in the nasal mucosa.

Rhinovirus and RSV are both capable of eliciting wheezing, but rhinovirus is more strongly associated with asthma exacerbations in children and adults^91^. Therefore, we hypothesized that the nasal cellular responses in rhinovirus participants with wheezing symptoms would be distinct from RSV participants with wheezing symptoms. To further understand cell subsets that may be associated with wheeze, we first evaluated the cellular composition of rhinovirus and RSV cases divided by wheeze group. We compared participants with rhinovirus or RSV who did not wheeze on exam and did not receive bronchodilators or steroids during their ED visit (No wheeze) with participants who had wheeze on exam and received bronchodilator and/or steroid therapy in the ED (Wheeze + treatment). Of note, swabs were obtained at any point during the ED stay and the specific time was not recorded relative to receipt of medical therapies. We noted that the rhinovirus wheeze+treatment cases were the most similar to each other compositionally, with greater variance in the other groups **(Figure 6G**). We applied a machine learning method to build a model to predict whether a rhinovirus sample belonged to the wheeze group and evaluated the importance of each cell subset in this prediction. Both immune and epithelial subsets were identified as important for the model, including subsets that were increased, decreased, or unchanged in wheezing participants **(Figure 6H**). A similar analysis in RSV cases revealed that some subsets were important for distinguishing wheeze for both viruses, including *XCL1/2+CTRAM*+ γδ T cells, neuronal cells, and *CXCL13+* CD8+ T cells, but that the majority of subsets identified for the RSV and rhinovirus wheezing models were specific to each virus **(Supplementary Figure 8E).** We found that multiple IFN-responsive cell subsets, including IFN-stim CD8+ cells, IFN-stim γδ T cells, and IFN-stim goblet cells, were decreased in frequency in rhinovirus wheeze cases, but not for RSV wheeze cases **(Figure 6I**). Furthermore, rhinovirus participants with wheezing symptoms had overall lower scores for IFNα and IFNγ **(Supplementary Figure 8F**). This may indicate that the bias toward an IL-13-responsive secretory state and away from an IFN-response epithelium observed in rhinovirus **(Figure 4**) is particularly pronounced in participants with wheeze.

As many participants in the wheeze group received systemic corticosteroids, we sought to determine the impact of corticosteroid treatment on nasal responses. In both rhinovirus and RSV wheeze group cases, epithelial and immune cells scored higher for a corticosteroid response signature **(Supplementary Table 5**)^92^ compared to the non-wheeze group **(Supplementary Figure 8G**). We compared cell subsets from participants from all viral groups who received corticosteroid treatment (systemic steroids in the ED or systemic or inhaled steroids in the week prior to presentation) with participants who did not. While IFN-stim goblet cells decreased in frequency in participants who had been treated with steroids, multiple cell subsets were enriched, including *BPIFA1+MUC5B+* goblet cells, mast cells, and *CXCL13+* CD8+ T cells **(Supplementary Figure 8H**). As *CXCL13* is typically expressed in CD4+ T peripheral helper cells (Tph), we confirmed that this subset expressed CD8+ T cell markers and was distinct from the *CXCL13*-expressing Tph subset also identified in this dataset **(Supplementary Figure 8I**). The *CXCL13*+ CD8+ subset was distinguished by *CXCL13* expression as well as upregulation of the steroid receptor *FKBP5*, suggesting that corticosteroid treatment may upregulate this chemokine in CD8+ T cells and alter B cell recruitment.

The steroid analysis suggests that some subset changes in wheeze groups may be due to corticosteroid treatment in the wheeze groups. However, cycling basal cells, *MUC5AC*+ deuterosomal cells and *CST1+* goblet cells were uniquely enriched in the rhinovirus wheeze cases **(Figure 6I**). Intriguingly, *CST1* was recently identified as a nasal transcriptomic marker of type 2-high asthma in adolescents^93^, is increased in induced sputum from adults with eosinophilic asthma^57^, and is upregulated in the airways of children with atopy^94^. We anticipated that *CST1*+ goblet cells express a transcriptional program important for type 2-polarized responses to rhinovirus. Indeed, the top three transcription factor activities were *STAT6*, *JUN*, and *AP1* **(Figure 6J**). *STAT6* is a master regulator of type 2 inflammation^95^, and *JUN* and *AP1* complexes regulate type 2 inflammation with a particularly important role in airway reactivity^96^. Furthermore, multiple markers of this cell subset (*CST1, ALOX15, TFF3, SERPINB3,* **Figure 6K**) are upregulated in polyps from individuals with chronic rhinosinusitis^73^. Together, our results strongly implicate *CST1+* goblet cells as an important cell state connecting rhinovirus responses to wheezing symptoms via local type 2 inflammation in the nose. Collectively, these findings highlight the capacity of this approach to discern phenotype-specific epithelial and immune programs *in vivo*, informing mechanistic hypotheses regarding the pathogenesis of severe RSV infection and rhinovirus-associated wheeze.

### Pediatric nasal cell atlas enables contextualization of genetic risk factors for childhood-onset asthma

Early childhood viral infections, particularly with rhinovirus or RSV, have been linked to the onset and later development of asthma^11^. At the population level, genome-wide association studies (GWAS) have identified numerous genetic risk loci associated with asthma; however, these studies do not indicate the specific cell types in which risk genes are active. This missing cellular context limits our understanding of how the combination of genetic risk and viral infection contributes to asthma development. We therefore asked whether our atlas of pediatric viral infection, integrated with GWAS data, could help address this gap. We applied the single-cell disease relevance score (scDRS) method, which scores individual cells based on their expression of genes implicated in GWAS studies^97^. For all cells in our dataset, we calculated the scDRS for childhood– and adult-onset asthma traits (COA and AOA, respectively), as well as height as a control trait. Squamous epithelial cells had the highest disease relevance scores for childhood-onset asthma, followed by goblet cells **(Figure 7A-B**). Interestingly, very few immune cells had significant enrichment of relevant genes for childhood-onset asthma in comparison with epithelial cells. In contrast, fewer epithelial cells but a greater number of immune cells had significant disease relevance scores for adult-onset asthma, with the highest scores in CD4+ T cells **(Figure 7A-B, Supplementary Figure 9A**). These results highlight the importance of the airway epithelium in childhood-onset asthma risk in comparison with adult-onset asthma risk.

**Figure 7:**
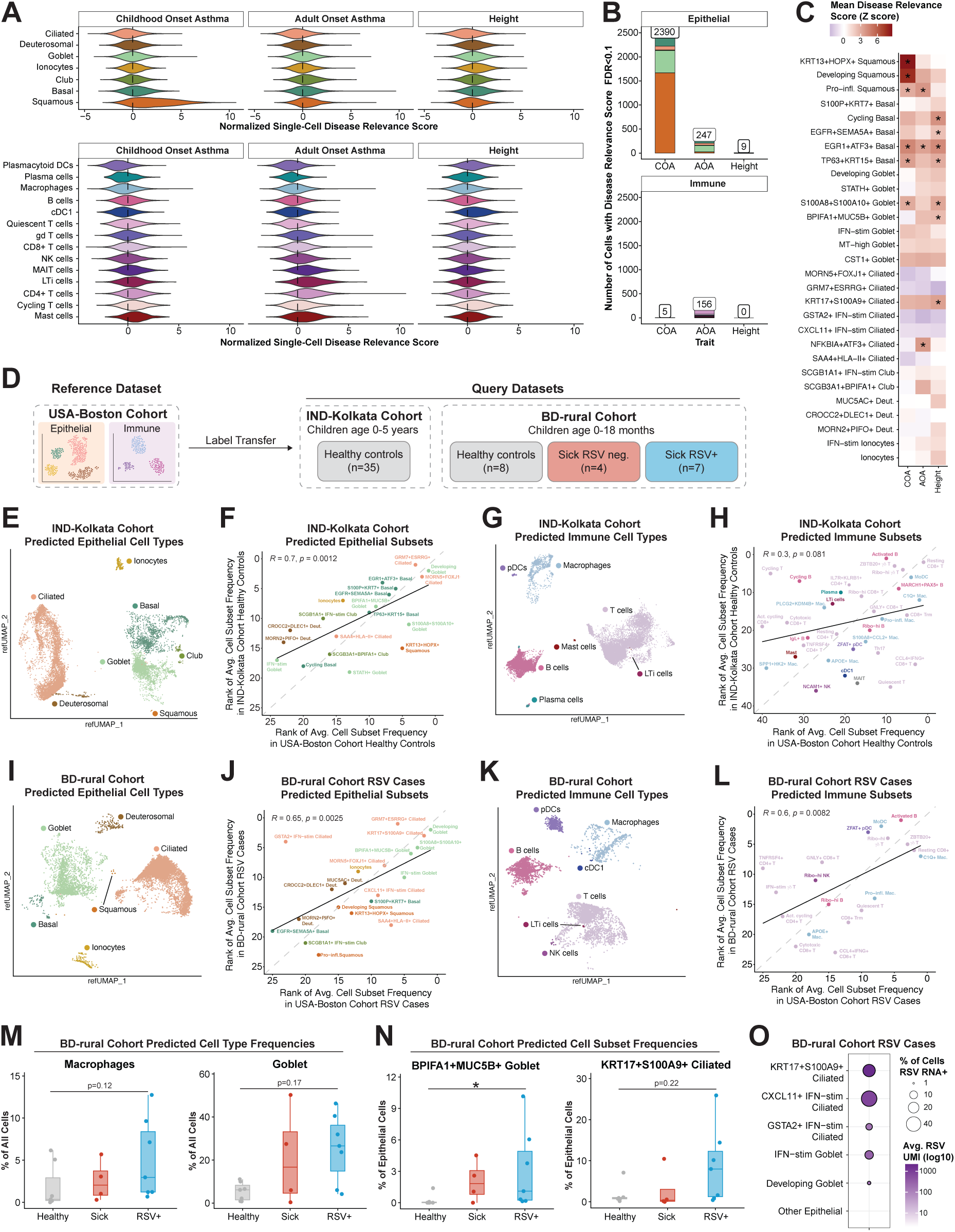
Applying the nasal cell atlas to understand genetic and geographical contributors to disease. **A.** Violin plot of normalized single-cell Disease Relevance Scores (scDRS) for childhood onset asthma, adult onset asthma, and height genome-wide association study (GWAS) traits, split by epithelial (top) and immune (bottom) cell type. **B.** Number of cells with a significant scDRS score at 10% FDR for childhood onset asthma (COA), adult onset asthma (AOA), and height. Cell types are colored as in **(A**). **C.** Cell type-trait association Z-score for childhood onset asthma (COA), adult onset asthma (AOA), and height across epithelial subsets. *FDR adjusted p<0.05. **D.** Schematic of label transfer approach. Cell type and cell subset annotations were transferred separately from epithelial and immune cells from the USA-Boston cohort to two separate query cohorts. See **Methods** and **Supplementary Table 2** for cohort details. **E,G**. UMAP of epithelial cells (n=13,514) **(E**) and immune cells (n=28,342) **(G**) from all samples in the IND-Kolkata cohort colored by predicted cell type. Cells with prediction score >0.8 are shown. **F, H.** Spearman correlation of cell subsets from controls ranked by average frequency among epithelial cells **(F**) and immune cells **(H**). X axis, rank in healthy controls in USA-Boston cohort. Y axis, rank in healthy controls in IND-Kolkata cohort. Subsets are colored by cell type of origin as in **E, G**. **I, K**. UMAP of epithelial cells (n=12,715) **(I**) and immune cells (n=5,266) **(K**) from all samples in the BD-rural cohort colored by predicted cell type. Cells with prediction score >0.8 are shown. **J,L.** Spearman correlation of cell subsets from RSV cases ranked by average frequency among epithelial cells **(J**) and immune cells **(L**). X axis, rank in RSV cases in USA-Boston cohort. Y axis, rank in RSV cases in BD-rural cohort. Subsets are colored by cell type of origin as in **I,K**. **M**. Frequency of select predicted cell types out of all cells per sample in BD-rural cohort, divided by disease group. Participants with <100 total cells were excluded. **N.** Frequency of select predicted epithelial cell subsets out of all epithelial cells per sample in BD-rural cohort, divided by disease group. Participants with <100 epithelial cells were excluded. **O.** Dotplot of RSV RNA+ cells in each epithelial subset in all RSV participants in BD-rural cohort. Dot size represents the percentage of cells in the subset that are viral RNA+. Color represents the average viral transcripts (UMI) across all cells in the subset. **M,N:** Boxplots represent the median (center line), interquartile range (box), and 1.5x the interquartile range (whiskers). Statistical test is Kruskal-Wallis test with Benjamini-Hochberg correction for multiple comparisons. Dunn’s post-hoc test, *p<0.05, **p<0.001, ***p<0.0001.

Next, we investigated the specific cell subsets which had the highest disease relevance scores. Among epithelial subsets, the score for childhood-onset asthma was strikingly enriched in *KRT13+HOPX+* squamous cells **(Figure 7C, Supplementary Figure 9B**). This subset expressed high levels of many cytokeratins (*KRT6A, KRT13, KRT78*), squamous markers (*SPRR2A*, *SPRR3*), and genes involved in epithelial remodeling or barrier integrity (*TMPRSS11E*, *PRSS27, CST6, ECM1*) **(Supplementary Figure 9C**). Notably, this same subset had the highest IL-13 response score **(Figure 4G**). The *KRT13+HOPX+* squamous cells in our dataset resemble “hillock” cells described as important progenitors in mouse and human airway injury^55,61^, suggesting that epithelial repair programs may overlap with asthma risk genes. To understand which genes were driving the disease relevance scores, we evaluated which genes had both a correlation with the disease relevance score and a positive gene weight, which serves as a correlate of GWAS risk. The genes contributing the most to the childhood onset asthma score in epithelial cells included many of the marker genes for the *KRT13+HOPX+* squamous subset **(Supplementary Figure 9D**), demonstrating the power of this approach to identify genes with individually weak contributions to genetic risk that when expressed in concert in a single cell may have important contributions to disease pathogenesis. Additionally, many T cell-relevant genes, including *ICOS, RORA,* and *CD247*, contributed the most to the adult-onset asthma score in immune cells. Together, these results demonstrate the utility of our dataset in contextualizing asthma GWAS data in terms of nasal cellular biology, and reveal that “hillock”-like squamous epithelial cells may be of particular importance in childhood onset asthma, consistent with a recent report^98^.

### Pediatric nasal cell atlas can be used as a reference for independent datasets

Geographic environment and genetic ancestry are known to contribute significantly to variation in cellular and tissue biology, leading to concerted efforts among human cell atlases to diversify the participants included. Additionally, while RSV infection is almost ubiquitous among children globally, the vast majority of childhood mortality from RSV takes place in low– and middle-income countries. Therefore, we sought to understand whether our atlas could be used as a viable reference in scenarios where the population, environment, and disease burden may be very different than in Boston, USA. To do this, we established independent participant cohorts in the area surrounding Kolkata, India (IND-Kolkata cohort, n=35, all healthy controls) and rural Bangladesh (BD-rural cohort, n=19, mix of controls and patients with respiratory infection) **(Figure 7D**). scRNA-seq library capture and sequencing was done locally using Honeycomb technology at each site, an important step in building capacity for future scRNA-seq studies without dedicated instrumentation.

First, we compared nasal cells from healthy children in the IND-Kolkata cohort (metadata provided in **Supplementary Table 2**) to the healthy controls in the USA-Boston cohort. After generating a dataset of 41,721 quality cells **(Supplementary Figure 10A-D**), we transferred labels separately from epithelial and immune cells. Predicted epithelial labels included the majority of expected cell types, including ciliated, goblet, basal, club, squamous, and deuterosomal cells, as well as ionocytes, and multiple cell subset labels were transferred for most cell types **(Figure 7E-F**). We also recovered both common (T cells, B cells, macrophages) and rare (plasma cells, mast cells, pDCs, LTi cells) immune cell types **(Figure 7G**), and many immune subsets were represented **(Figure 7G-H**). The majority of cells in our dataset had prediction scores close to 1, though the distribution was slightly wider in immune cells and varied by cell type **(Supplementary Figure 10E-F**). Analyzing the frequency of predicted cell types across the individual samples revealed consistent detection of both epithelial and immune cell types **(Supplementary Figure 10G**). To compare the two cohorts on the cell subset level, we ranked the abundance of epithelial and immune subsets from each cohort **(Figure 7F, 7H**). Healthy controls in the IND-Kolkata cohort had high frequencies of *GRM7+ESRRG+* ciliated and *MORN5+PIFO+*ciliated cells as well as multiple basal cell subsets **(Figure 7F**), similar to the controls in the USA-Boston cohort **(Figure 1I**, **Figure 3F-G**). There was less concordance in frequency of immune subsets, though quiescent populations such as resting CD8+ T cells, ribo-hi γδ T cells, and *MARCH5+PAX5+* B cells ranked highly in frequency in both cohorts **(Figure 7H**). Together, these analyses indicate that our atlas can be used as a reference to find shared epithelial and immune phenotypes among healthy children from different environments.

Next, we evaluated similarities in infection responses between children in the BD-rural cohort and the virally-infected cases in the USA-Boston cohort. The BD-rural cohort was divided into children who had no respiratory symptoms (Healthy, n=8), had respiratory symptoms but were negative for RSV (Sick, n=4), and symptomatic RSV+ cases (n=7) **(Figure 7D, Supplementary Table 2**). We generated a dataset of 17,981 high-quality cells **(Supplementary Figure 10H-K**) with single-cell detection of RSV RNA in 4/7 RSV samples **(Supplementary Figure 10L**). We used the same label transfer approach to annotate epithelial and immune cells, and similarly found recovery of expected cell types, significant correlations between subset rankings, and strong prediction scores **(Figure 7I-L, Supplementary Figure 10M-N**). Notably, *KRT17+S100A9+* ciliated cells were among the most frequent epithelial subsets in both the BD-rural RSV cases and USA-Boston RSV cases **(Figure 7J**). Concordance in immune subset frequencies was improved when transferring between infected cohorts **(Figure 7L**), indicating that the broader immune heterogeneity present during infection may facilitate more robust label transfer.

Investigation of cell type frequencies between groups in the BD-rural cohort revealed trends of enriched macrophages and goblet cells in RSV cases **(Figure 7M**), as well as specific increases in *BPIFA1+MUC5B+* goblet cells and a trending increase *KRT17+S100A9+* ciliated cells in RSV cases **(Figure 7N**), consistent with our previous results. Finally, we evaluated which of the predicted epithelial subsets contained RSV RNA. Strikingly, we observe RSV RNA in the same epithelial subsets as in the USA-Boston cohort, including *KRT17+S100A9+* ciliated cells and *CXCL11*+IFN-stim ciliated cells, as well as in a few other subsets **(Figure 7O, Supplementary Figure 10O**). Together, these results confirm the ability of our atlas to serve as a reference for both healthy and infected datasets, and highlight the finding of RSV-specific *KRT17*+ ciliated cells as a conserved phenotype across cohorts.

## Discussion

Here, we present a comprehensive atlas of nasal epithelial and immune cell responses to three common viral infections in young children. We sought to define how SARS-CoV-2, rhinovirus, and RSV shape the cellular landscape of the pediatric nasal mucosa and how virus-specific cell states relate to clinical outcomes. By systematically profiling epithelial and immune compartments, we identified distinct infected epithelial cell states in SARS-CoV-2 and RSV and delineated broader convergent and divergent changes in nasal cellular composition across infections. SARS-CoV-2 infection induced a robust IFN response, whereas rhinovirus infection instead drove a secretory epithelial program marked by IL-13-responsive gene expression, and RSV elicited heightened cytotoxic and inflammatory immune activity. We found that severe RSV cases exhibited increased type I and type II IFN-responsive programs, while rhinovirus-associated wheezing showed enrichment of goblet and deuterosomal cell subsets. By interpreting asthma GWAS data through the lens of our atlas, we identified a hillock-like squamous epithelial subset with particular relevance to childhood-onset asthma. Finally, we demonstrated that this atlas serves as a high-resolution reference for annotating independent pediatric nasal datasets and uncovering shared cellular states. Together, these findings suggest that common respiratory viruses engage distinct epithelial response programs in the pediatric nasal mucosa, which in turn correspond to divergent immune activation patterns and clinical outcomes.

SARS-CoV-2 infection is typically mild in young children, yet its prevalence and continued circulation underscores the importance of defining pediatric mucosal responses to this virus. By mapping viral transcripts to individual cells, we observed that the majority of SARS-CoV-2 RNA in the pediatric nasal mucosa was found in a ciliated epithelial subset marked by strong NF-κB–responsive gene expression. This transcriptional state closely mirrors a SARS-CoV-2 RNA-high epithelial subset previously identified in the adult nasal mucosa as consistently enriched across SARS-CoV-2 viral variants^19^, as well as a “hyper-infected” ciliated epithelial population described in an adult human SARS-CoV-2 challenge study^18^. Shared features of these populations include induction of canonical NF-κB feedback regulators (*NFKBIA, NFKBIZ, TNFAIP3*), AP-1-associated immediate-early genes indicative of epithelial activation (*ATF3, EGR1, FOS, JUN*), and markers of cellular stress and metabolic adaptation (*DDIT3, DNAJB1, GDF15*). NF-κB-driven transcriptional programs are functionally required for efficient SARS-CoV-2 replication in epithelial cell lines and nasal organoid models^99,100^. Importantly, comparison with rhinovirus and RSV infections demonstrated that this epithelial activation state is specific to SARS-CoV-2 in the pediatric nasal mucosa. Together, these findings suggest that a conserved NF-κB-responsive epithelial program may represent a relevant target for SARS-CoV-2 antivirals.

Although the SARS-CoV-2-infected epithelial cell subset is not primarily defined by a canonical interferon-stimulated gene program, it expresses *IFNL*, which induces a robust interferon response in surrounding epithelial cells in mucosal tissues^101,102^. Consistent with this, we observed increased IFNα response signature scores in epithelial cells and enrichment of multiple interferon-stimulated cell subsets – including goblet, club, ciliated epithelial cells and T cells – in SARS-CoV-2 cases compared to rhinovirus and RSV. Prior scRNA-seq studies comparing adult and pediatric SARS-CoV-2 infection have similarly reported heightened interferon responses and higher baseline expression of interferon-stimulated genes in children^25,26,28^, and it has been proposed that this contributes to the generally mild clinical course of SARS-CoV-2 infection in children^103^. Our data are consistent with this model and further suggest that this pronounced interferon landscape distinguishes SARS-CoV-2 from rhinovirus and RSV infection in early childhood.

In contrast to SARS-CoV-2, rhinovirus infection was characterized by a diminished interferon response, loss of ciliated epithelial cells, and a shift toward secretory and goblet cell differentiation accompanied by induction of IL-13-responsive genes. Consistent with these findings, prior studies in both clinical samples and experimental infection models have reported reduced expression of ciliary genes and increased goblet cell–associated programs during rhinovirus infection^104–107^. Recent work further demonstrated that epithelial interferon signaling is critical for restricting rhinovirus replication, and that disruption of this response permits NF-κB-, NLRP1-, and IL-1β-driven inflammatory signaling and mucus production^33^. Given that ciliated epithelial cells are a primary target of rhinovirus infection^108,109^, the pronounced reduction in ciliated cells we observed may reflect preferential infection and subsequent sloughing of ciliated cells. In line with this model, we identified multiple secretory cell subsets enriched in rhinovirus cases, including a *MUC5AC+* deuterosomal population. Deuterosomal cells are thought to represent transient intermediates in multiciliated cell differentiation^53^; however, their enrichment in rhinovirus infection may reflect altered epithelial differentiation trajectories with potential diversification toward secretory or ciliated fates. Together, these findings suggest that, *in vivo*, rhinovirus infection preferentially engages epithelial differentiation pathways associated with mucus production downstream of Type 2 cytokine signaling, rather than an interferon-mediated antiviral response.

The epithelial response to RSV infection *in vivo* remains poorly understood. RSV primarily infects ciliated cells *in vitro*, although productive infection of basal cells has also been described^110,111^. In our *in vivo* dataset, the epithelial subset harboring the majority of RSV RNA exhibited a striking transcriptional profile characterized by programs related to squamous-like activation (*KRT17, KRT23, SPRR1B*), wound repair and extracellular matrix remodeling (*LAMB3*, *PLAUR*, *TIMP2*), and inflammation (*S100A9, C3, CXCL8*) alongside remarkably low expression of ciliated markers and ISGs. RNA velocity analysis suggested that this subset most likely arose from ciliated cells, consistent with viral injury-induced reprogramming following infection. Early work demonstrated that RSV infection induces NF-kB dependent expression of cytokeratin 17 (KRT17) in respiratory epithelial cells, with localization to sites of syncytia formation^112^, providing initial experimental evidence that RSV can drive cytoskeletal remodeling through inflammatory signaling. Our findings extended these observations to an *in vivo* context, and showed that this *KRT17+S100A9+* activated epithelial state is highly specific to RSV-infected cells. KRT17 has also been linked to aberrant epithelial activation and dysregulated wound repair in other respiratory diseases, including pro-fibrotic epithelial programs in idiopathic pulmonary fibrosis^113,114^. Together, these observations suggest that this RSV-associated epithelial program reflects a cell intrinsic response to viral infection, and we hypothesize that this state may promote viral persistence and dampen effective intrinsic antiviral responses.

In addition to this unusual infected epithelial cell state, RSV cases exhibited a heightened inflammatory and cytotoxic immune profile. Cell-cell communication analyses revealed macrophage-centered signaling, and network-level analyses identified a coordinated circuit linking RSV-infected epithelial cells with NK cell, T cell, and macrophage subsets. Several RSV-enriched subsets, including *KRT17+S100A9+* ciliated cells, activated cycling CD4+ T cells, and cytotoxic CD8+ T cells, correlated with clinical outcomes such as illness duration, need for respiratory support, and hospital admission across all patients in the cohort. Consistent with this observation, profiling in the blood and airway has demonstrated innate immune remodeling in severe RSV, including monocyte activation, altered NK cell abundance and function, and IL-36 associated neutrophilic inflammation^24,40–42,87,88,115^. In our cohort, RSV participants admitted to the ICU displayed a cellular composition driven by IFN-stimulated epithelial and immune subsets, along with elevated type I and type II IFN response signatures and higher nasal viral loads. In contrast, prior studies have generally reported diminished airway IFN responses in severe or intubated RSV cases^39,42,86,116,117^, despite associations between nasal viral load and IFN signaling^118,119^. These differences likely reflect variation in disease stage and sampling site, as our cohort was sampled in the emergency department and may represent an earlier phase of illness. Together, these findings support a model in which RSV elicits a comparatively limited epithelial-intrinsic antiviral program while driving robust innate and cytotoxic immune activation, in contrast to SARS-CoV-2, where a stronger epithelial interferon response may help restrict viral spread within the mucosa. In this heightened inflammatory context, *KRT17*+ epithelial remodeling and elevated IFN signaling may reinforce inflammatory circuits, contributing to immune-mediated pathology rather than effective viral containment.

Our dataset also enabled identification of cellular subsets associated with virus-induced wheeze. Some of these phenotype-specific states, including mast cells and *CXCL13*+ CD8+ T cells, may in part reflect transcriptional responses to therapeutic steroids, and disentangling treatment effects from disease biology will require further study. Clinically, rhinovirus is more strongly linked to subsequent asthma exacerbations, and rhinovirus-induced wheeze more often reflects an asthma-like, type 2–skewed inflammatory phenotype that responds to standard asthma-directed therapies, whereas RSV-associated wheeze typically arises from bronchiolitis-driven inflammation and does not demonstrate the same treatment responsiveness^6,11,91^. In line with this, wheeze-associated signatures in our cohort differed markedly between rhinovirus and RSV cases. Rhinovirus-associated wheeze was characterized by further polarization toward a secretory, IL-13-responsive epithelial program with reduced IFN-responsive signatures. These findings suggest that robust interferon responses may mitigate wheezing severity, while type 2-biased immunity may predispose to asthma-like symptoms. We identified a *CST1*+ goblet cell subset enriched specifically in rhinovirus-induced wheeze, with features consistent with STAT6-mediated transcriptional regulation downstream of IL-4 and IL-13 signaling^120^. CST1 expression has been associated with eosinophilic asthma and correlates with Th2 cytokine activity in adults^57,58,60^. The presence of this epithelial state in young children following infection raises the possibility that type 2-skewed disease trajectories may begin early in life. It remains to be further characterized whether rhinovirus drives such disease pathologies and subsequent risk for asthma or if rhinovirus-associated wheezing in young children is a marker of susceptibility for a Th2-skewed response.

By interrogating which specific cell subsets in the pediatric nasal mucosa are enriched for genetic risk of asthma onset, we demonstrated the broader utility of this atlas beyond acute infection responses. Genetic risk for asthma intersects with susceptibility to rhinovirus-induced wheeze^7,9^, and rhinovirus infection has been shown to upregulate childhood-onset asthma-associated genes, particularly in non-ciliated epithelial cells^121^. In our dataset, childhood-onset asthma GWAS enrichment scores were highest in squamous epithelial cells, and most pronounced in a *KRT13+HOPX+* squamous subset. This population resembles “hillock” cells, an epithelial subset implicated in airway repair and remodeling^50,55,61,122^. Emerging evidence further links KRT13+ hillock-like epithelial states and squamous differentiation programs to childhood-onset asthma, pediatric wheeze, and allergic airway inflammation^36,98,123^. Together, these findings suggest that infection-induced activation or expansion of this squamous repair program may intersect with inherited asthma risk and shape long-term airway remodeling trajectories.

Finally, we demonstrated that this dataset serves as a robust resource for annotating independent nasal scRNA-seq samples. Using our atlas as a reference, we successfully transferred annotations to both healthy and infected pediatric nasal samples generated in India and Bangladesh using a more portable technology, with strong concordance in both cell type identity and relative subset abundances across sites. Notably, *KRT17+S100A9+* ciliated cells identified in the BD-rural cohort similarly harbored RSV RNA, reinforcing the reproducibility of key epithelial states across geographic contexts. To our knowledge, this represents the first published single-cell RNA-sequencing dataset generated locally in Bangladesh, and one of the few locally produced pediatric airway scRNA-seq datasets from low– and middle-income countries. This work illustrates a scalable model in which deep reference profiling at one site can support streamlined, lower-depth profiling across additional settings, enabling greater flexibility in technology selection and study design. Advancing our understanding of nasal biology and respiratory infection requires inclusion of diverse geographies, ancestries, and populations, particularly given that the burden of childhood respiratory disease is greatest in low– and middle-income countries^3,5,43,44,46^. By prioritizing partnership and capacity building, we aimed to establish a sustainable framework for future studies. Moving forward, deeper integration of datasets across locations will allow us to assess both shared and region-specific cellular features of pediatric respiratory disease.

In summary, we establish a comprehensive single-cell atlas of the pediatric nasal mucosa during common respiratory viral infections, defining virus-specific epithelial and immune cell states and linking them to clinical outcomes. Beyond characterizing acute infection responses, this atlas enables hypothesis generation regarding mechanisms of disease severity and asthma risk, and provides a reference framework for annotation and comparison of future pediatric nasal cohorts. By integrating viral detection, cellular phenotypes, and genetic risk, our findings offer a foundation for understanding how early-life respiratory infections may influence longer-term airway biology. Future longitudinal studies and integration with functional experiments will be necessary to define the temporal dynamics and causal relationships underlying these responses across diverse populations.

### Limitations of the study

This study has several limitations. First, each participant was sampled at a single time point, limiting our ability to assess the temporal dynamics of viral replication and host response. Children were sampled at presentation to the emergency department, representing different stages of illness, and illness duration was self-reported by the participant’s parents. Second, our focus on early childhood restricts our ability to compare infants and toddlers directly or to fully evaluate age-dependent differences within the cohort. Third, cell numbers were limited in some samples, and certain cell types such as neutrophils were not detected, likely due to freeze-thaw processing. Fourth, single-cell clustering and annotation are inherently influenced by upstream analytical choices and remain partially subjective; we have therefore provided detailed rationale and methodological transparency to support reproducibility. Finally, detection of viral RNA within a cell does not definitively establish productive infection and should be interpreted as evidence of viral transcript presence rather than replication competence. In contrast to RSV and SARS-CoV-2, we detected rhinovirus transcripts only rarely in our single-cell data. This limited detection may reflect rapid sloughing or death of infected ciliated epithelial cells, lower levels or shorter duration of viral RNA persistence, or the substantial genomic diversity of rhinoviruses^91^, which may reduce alignment efficiency to reference genomes. As a result, we were not able to define rhinovirus-infected epithelial cell states with the same resolution as for RSV and SARS-CoV-2. Dedicated approaches, including targeted viral enrichment or earlier sampling relative to symptom onset, will likely be required to more comprehensively characterize rhinovirus-infected epithelial populations *in vivo*.

## Acknowledgements

We thank the study participants and their families for enabling this research, the clinical support staff at each site for their efforts in sample collection, and members of the Ordovas-Montanes and Horwitz labs for thoughtful discussion and feedback. From the BCH ED, we thank Tom Stivers for protocol management and oversight, and clinical research coordinators John Schultz, Michelle Du, Rebecca Wolf, Joseph Griffiths, Deirdre Flanagan, Abigail Bryant, Carolyn Drescher, Joseph Kanaan, and Aniyah Travis for patient recruitment and sample collection. For support of sample collection and data generation for the BD-rural cohort, we thank Naito Kanon, Himadree Sarkar, Mohammad Shameem Hassan, Shakiul Kabir and the Mirzapur Field Team. We thank the nurses and technicians of the Institute of Child Health, Kolkata, for help with recruitment of the children and collection of swabs from them. We thank all members of the BCH Cell Discovery Network for computational support and analytical insight. We also thank Maria Gutierrez-Arcelus and Vitor Rezende Da Costa Aguiar for helpful discussions related to scDRS analysis. We thank Erica Rutherford for support in data curation. We sincerely thank Arlene Sharpe and all members of the Sharpe laboratory for insightful discussion and feedback as well. We thank Juliane Weller for making the mg2sc pipeline for kraken2 single-cell analysis publically available. This has been made possible in part by CZI grant DAF2021-237665 and grant DOI https://doi.org/10.37921/881723wbyqpp from the Chan Zuckerberg Initiative DAF, an advised fund of Silicon Valley Community Foundation (funder DOI 10.13039/100014989), awarded to AKShalek, SCG, TIS, AKSesay, CSQB, LMY, PM, SS, BHH, and JOM, as well as a Grant DAF2023-330879 to JOM and Grant DAF 2023-330382 to JMLW. JOM is a New York Stem Cell Foundation – Robertson Investigator. JOM was supported by the AbbVie-Harvard Medical School Alliance, the Leona M. and Harry B. Helmsley Charitable Trust, The Pew Charitable Trusts Biomedical Scholars, The Broad Next Generation Award, The Chan Zuckerberg Initiative Pediatric Networks, The Mathers Foundation, The Kenneth Rainin Foundation, The Massachusetts Consortium for Pathogen Readiness, NIH R01 HL162642, NIH R01 DE031928, NIH P30 DK034854, the NIH CIDER Network, Project NextGen, and The Cell Discovery Network, a collaborative funded by The Manton Foundation and The Warren Alpert Foundation at Boston Children’s Hospital. JMLW was supported by NIH Training Grant 5TL1TR002543. LJJ was supported by NIH 5T32HD098061 Neonatal Research Training Program to Boston Children’s Hospital, the Marshall Klaus Perinatal Research Award, a Thrasher Foundation Early Career Award, and a Gerber Foundation Novice Award.

## Author contributions

Contributions are listed according to CRediT criteria: https://credit.niso.org/

Conceptualization: JMLW, LJJ, YT, AC, AKShalek, PM, YH, SS, BHH, JOM;

Data curation: JMLW, LJJ, YT, AC, FT, CML, PCD, LT, AM, SG, ASK, YH,

Formal analysis: JMLW, LJJ, MEB, HRM;

Funding acquisition: AKShalek, SCG, TIS, AKSesay, CSQB, WG, LMY, PM, SS, BHH, JOM;

Investigation: JMLW, LJJ, YT, AC, EL, AF, FT, DPK, PCD, ARM, LT, AMT, AM;

Methodology: JOM, JMLW, LJJ, BHH, YT, SS, YH, PM, AC;

Project administration: JMLW, LJJ, YT, AC, AKShalek, PM, YH, SS, BHH, JOM;

Resources: YT, AC, AKShalek, PM, YH, SS, BHH, JOM;

Software: JMLW, LJJ, MEB, HRM, CML, PCD, SG

Supervision: JMLW, LJJ, PM, YH, SS, BHH, JOM;

Validation: JMLW, LJJ;

Visualization: JMLW, LJJ, MEB, KK;

Writing-original draft: JMLW, LJJ, JOM;

Writing-review & editing: JMLW, LJJ, YT, AC, AKShalek, PM, YH, SS, BHH, JOM

## Methods

### Study Cohort Details

Participants were recruited from 4 sites as described below. Participant metadata for BCH and MGH cohorts is summarized in **Supplementary Table 1**, and information for all individual participants is available in **Supplementary Table 2**.

### USA-Boston cohort: BCH

A convenience sample of children under 5 years of age presenting to the Emergency Department (ED) at Boston Children’s Hospital (BCH) with either 1) signs of viral respiratory illness including cough, nasal congestion, wheezing or difficulty breathing, or 2) minor traumatic injuries including minor head trauma and simple lacerations without symptoms of respiratory viral infection within the last 30 days were identified by screening of the electronic health record. Following informed consent from a parent or legal guardian, a research nasopharyngeal (NP) swab was obtained from each child. When available, discarded viral transport media from clinically indicated nasal sampling were also collected. In addition, parents were administered a questionnaire regarding past medical history, current medication usage, and current symptomatology. Following patient disposition, review of the electronic health record was performed to obtain patient clinical parameters related to the patients visit including vital signs, prescribed medications, ED disposition, and hospital length of stay, as well as demographic data. All clinical and demographic data was entered into a REDCap database. This study was approved by the Boston Children’s Institutional Review Board under protocol number P00028229.

Children who presented with symptoms consistent with respiratory viral infection were first assigned to one of the viral groups (SARS-CoV-2, RSV, or rhinovirus) based on detection of one of these viruses by clinically indicated testing and/or by qPCR performed on viral transport media. Children who presented with minor traumatic injuries were assigned to the control group. For 126/132 samples, when processing the research NP swabs for single-cell RNA-sequencing, we saved supernatants at multiple steps, combined and stored them for each sample, and repeated the qPCR to confirm infection status (see assay details below). Any control samples with a positive value for SARS-CoV-2, RSV-A, RSV-B, rhinovirus/enterovirus, influenza A, or influenza B were re-classified as asymptomatic cases (n=5). Any samples in the viral groups with positive values for more than one of the tested viruses were re-classified as coinfection cases (n=7). Asymptomatic and coinfection cases were included in cell clustering and annotation, but excluded from virus-specific comparisons.

Participants who were admitted to the hospital from the ED were categorized as either admitted to a standard pediatric inpatient unit (Floor) or admitted to the Intermediate Care Program, Medical Intensive Care Unit, or Medical-Surgical Intensive Care Unit (categorized together as ICU) at BCH. Groups of patients who were wheezing were defined by the following characteristics: (1) presence or absence wheeze documented on physical exam in ED physician note; (2) receipt of wheezing-directed therapy in the ED, including nebulized albuterol, IV magnesium sulfate, and/or dexamethasone. Participants were categorized in the “No wheeze” group if they did not have documented wheeze and did not receive wheezing-directed therapies in the ED. Participants were categorized in the “Wheeze + treatment” group if they had documented wheeze and received wheezing-directed therapies in the ED.

Illness severity was quantified using a modified severity score based on the Bronchiolitis Score of Sant Joan de Deu (BROSJOD)^48^. We included the age-specific parameters for heart rate and respiratory rate, oxygen saturation, and accessory muscle use/retractions, all according to the BROSJOD criteria. We used the first recorded vital signs, obtained prior to medical therapy. We did not include the wheeze/rates or air entry components of the BROSJOD score as these were rarely reported in the BCH electronic medical record (EMR).

### USA-Boston cohort: MGH

Healthy children aged from 1 month to 5 years receiving care at Massachusetts General Hospital (MGH) Chelsea Healthcare Center were enrolled during their visit under an MGH Institutional Review Board-approved study. Written informed consent was acquired from parents/legal guardians. Demographic and clinical variables were extracted from participants’ EMR.

### IND-Kolkata cohort

Parental consent was obtained after providing adequate information about the research to obtain a nasal swab from children between 6 months and 5 years. None of the children was suffering from any infection. Most children had come for regular physical check-up or for immunization; swabs were collected before immunization. All sample collection procedures were minimally invasive and well tolerated, and no adverse events were reported. Ethical approvals for sample collection were provided by the Institutional Ethics Committees of the Institute of Child Health, Kolkata, and the National Institute of Biomedical Genomics, Kalyani, India.

### BD-rural cohort

Eligible participants were recruited in a rural community located in the Mirzapur subdistrict in Tangail district, Bangladesh (BD-rural), where Child Health Research Foundation (CHRF) has maintained a demographic-based surveillance system since 2007^124^. Written informed consent was obtained from the parents or legal guardians of all participants prior to sample collection. Enrolled participants were children less than 2 years of age who were identified by village health workers as sick or healthy. A child was considered sick if they met the WHO inclusion criteria for RSV community surveillance (defined as having at least one of the following: shortness of breath, cough, sore throat, coryza)^125^. Age, sex and location-matched healthy participants were recruited as the control group; these were children who did not meet the RSV inclusion criteria, did not have any other illness at the time of recruitment and did not report use of any medications. Samples were collected during two time periods: October to November 2023 and September to October 2024. The Ethical Review Committee of the Bangladesh Shishu Hospital and Institute approved the study (Admin/BSHI/2023/923/1).

## Method Details

### Resource availability

Further information and requests for resources and reagents should be directed to and will be fulfilled by the lead contact, Dr. Jose Ordovas-Montanes.

### Sample collection

Nasopharyngeal swabs were collected by a trained healthcare provider using FLOQSwabs (Copan). The swab was placed immediately into a cryogenic vial containing 2mL of heat inactivated fetal bovine serum (FBS, Corning) with 10% dimethyl sulfoxide (DMSO, Sigma). Cryovials were then placed into a room temperature Mr. Frosty Freezing Container (Thermo Fisher Scientific), and stored in a –80C freezer overnight. Samples were moved to individual boxes and kept at –80C.

### Dissociation of single cells from nasal swabs

Swabs in freezing medium (90% FBS, 10% dimethylsulfoxide) were stored at –80C until immediately before dissociation. A detailed sample protocol can be found here: https://www.protocols.io/view/czi-pediatric-nasopharyngeal-swab-processing-for-1-6qpvr49jogmk/v1. Briefly, nasal swabs in freezing medium were thawed, and each swab was rinsed with RPMI (Thermo Fisher Scientific) before incubation in 1 ml RPMI, 10 mM DTT (Sigma) for 15 min at 37 °C with agitation. Next, the nasal swab was incubated in 1 ml Accutase (Sigma) for 30 min at 37 °C with agitation. The 1 ml of RPMI with 10 mM DTT from the nasal swab incubation was centrifuged at 400g for 5 min at 4 °C to pellet cells, the supernatant was saved, and the cell pellet was resuspended in 1 ml Accutase and incubated for 30 min at 37 °C with agitation. The original cryovial containing the freezing medium and the original swab washes were combined and centrifuged at 400g for 5 min at 4 °C. The supernatant was saved and the cell pellet was then resuspended in RPMI with 10 mM DTT and incubated for 15 min at 37 °C with agitation and centrifuged as above. The cell pellet was then resuspended in 1 ml Accutase and incubated for 30 min at 37°C with agitation. All cells were combined following Accutase digestion and filtered using a 40-μm nylon strainer. The filter and the swab were washed with RPMI with 10% FBS and 4 mM EDTA, and all washes were combined. Dissociated, filtered cells were centrifuged at 400g for 10 min at 4 °C and resuspended in 200 μl RPMI with 10% FBS for counting. 10,000 live cells from each sample were combined into pools of 3-4 samples. The pooled cells were centrifuged at 400g for 5 min at 4C, then resuspended in 500uL PBS+1% BSA and centrifuged again as above. The pellet was resuspended in 43.3 uL of PBS+1% BSA to proceed to scRNA-seq. 58/132 samples were additionally subjected to live cell enrichment on the LeviCell 1.0 instrument. After counting, cells were pooled at equal ratios. The pooled cells were centrifuged at 300g for 5 minutes and resuspended in 230 uL. of 150mM Levitation Agent (Levitas Bio) diluted in RPMI+10% FBS. The standard cell enrichment protocol was run. After collecting the enriched cell population, cell counting was repeated. 10,000 live cells times the number of samples in the pool were transferred to a new tube, washed and resuspended in PBS+1% BSA as above to proceed to scRNA-seq.

### USA-Boston cohort: Single-cell RNA-seq

For participants in the USA-Boston cohort, pooled samples were processed using the Chromium NextGEM Single Cell 3’ Kit v3.1 with dual indices (10X Genomics) per the manufacturer’s instructions. Library quality was evaluated using the Agilent TapeStation 4200 (Agilent) and concentration was measured using the Qubit 1xdsDNA High-Sensitivity Assay Kit (Qubit). Sequencing was performed on the NovaSeq 6000 (Illumina) at the Broad Institute Sequencing Core with a targeted read depth of 25,000 reads per cell: read 1, 28; read 2; 90, index 1, 10; index 2, 10.

### IND & BD cohorts: Single-cell RNA-seq

For participants in the IND-Kolkata and BD-rural cohorts, cells were isolated using the same dissociation protocol as above. scRNA-seq libraries were generated for 10,000 cells for each individual sample using the HIVE™ scRNAseq v1 Processing Kit (Honeycomb Biotechnologies) or the HIVE CLX scRNAseq Kit (Honeycomb Biotechnologies) and prepared according to the manufacturer’s protocol. Libraries were sequenced on NextSeq 2000 or NovaSeq 6000 platforms (Illumina) with a targeted read depth of 25,000 reads per cell: read 1, 25; read 2, 50; index 1, 8; index 2, 8.

### USA-Boston cohort: Bulk RNA-seq

Aliquots of 5,000 cells from each sample were transferred to new tubes. These pellets were spun down at 400g for 5 min at 4C and resuspended in 50uL of RLT buffer (Qiagen) + 1% 2-mercaptoethanol (BME, Sigma). The lysed cells were snap frozen and stored at –80C. Libraries were prepared using the previously described SmartSEQ2 protocol^126^. Briefly, RNA was purified from 50 µL of cell lysate using 2.0X SPRIselect beads (Beckman-Coulter). Purified RNA was reverse transcribed into cDNA and whole transcriptome amplification was carried out. Libraries were fragmented and dual indexed using the Nextera XT Library Prep Kits (Illumina). Sequencing was performed on the NovaSeq 6000 or NextSeq 2000 at the Broad Institute Sequencing Core with a targeted read depth of 10 million reads per sample, read 1, 59; read 2; 59, index 1, 10; index 2, 10.

### USA-Boston cohort: Viral qRT-PCR

1mL RLT+BME was added to the combined supernatants collected during research NP swab processing to lyse any remaining cells. This lysate was then aliquoted into 1mL/cryovial, snap frozen, and stored at –80C until use. qRT-PCR was done on both scRNA-seq swab supernatants to confirm viral infection and on clinical swab samples to preliminarily assign groups. For both types of samples, RNA was extracted using QIAamp MinElute Virus Spin Kit (Qiagen) following the manufacturer’s instructions. 1-step standard RT-qPCR was performed on QuantStudio™ 7 Flex Real-Time PCR System (Thermo Fisher Scientific) using TaqMan assays to identify viral infections. Supernatants from 126/132 participants were tested for B-ꞵCoV E gene, SARS-CoV-2 S gene, RSV-A, RSV-B, influenza A, influenza B, rhinovirus/enterovirus, and RPP30 as the internal control. All samples were negative for influenza A and B. Samples with a cycle threshold (CT) value <40 were considered positive. All CT values reported in figures refer to scRNA-seq swab PCR.

### BD cohort: RSV Detection

RNA was extracted from 200 µL of the specimen using Quick RNA Viral kit (Zymo Research) following the manufacturer’s protocol. qPCR for RSV detection and subgrouping was performed using 6.25 µL RNA with Luna Universal Probe One-Step RT-qPCR Kit (New England Biolabs) and previously published RSV-specific primers and probes^127^. PCR cycling conditions were 50°C for 30 minutes, 95°C for 5 minutes, and 40 cycles of 95°C for 15 seconds and 60°C for 30 seconds. A true sigmoidal amplification curve of ct-value <35 confirmed RSV positivity.

### Quantification & Statistical Analysis

#### Sample demultiplexing

Bulk RNA-seq libraries were demultiplexed using Illumina bcl2fastq (v2.20). SNPs specific to each participant were called using Genome Analysis Toolkit (GATK4) workflows provided by the Broad Institute using default settings^128^. Paired FASTQs were converted into unmapped BAM files and aligned to the GRCh38 genome. Duplicate marking and base recalibration was performed and haplotype caller was used to generate a variant call file (VCF) for each participant. A merged VCF file was then created by aggregating the VCFs from all participants in each pool.

SNP-based genetic demultiplexing of pooled scRNA-seq samples was performed as previously described^129–131^. Briefly, scRNA-seq pools were initially demultiplexed in a genotype-agonistic method through the Freemuxlet variation of the Popscle package. Freemuxlet performs unsupervised clustering of single cells based on common SNPs identified within transcripts and assigns each cell as a nameless donor or doublet^130^. In addition to assigning cells, a VCF is generated for identifying SNPs of each donor. These VCFs are then compared to the aggregated VCF from each pool and matched based on genotype similarity (vcf-match-sample-ids)^131^. Freemuxlet and participant labels were then added to each cell. Cells identified as inter-participant doublets were excluded from further analysis.

#### scRNA-seq pre-processing

To detect both host and viral transcripts, we built a custom reference genome combining human GRCh38-2020-A (Ensembl 93), SARS-CoV-2^66^, RSV-A (GenBank: KY654518.1)^132^, RSV-B (GenBank: MZ516105.1)^133^, Rhinovirus C NAT001 (GenBank: EF077279.1)^69^, Rhinovirus B104 (GenBank: FJ445137.1)^134^, Rhinovirus B48 (GenBank: DQ473488)^69^, and Enterovirus E19 (GenBank: AY302544.1)^135^. For rhinovirus/enterovirus genomes, these genomes were selected for the custom reference genome based on Kraken2^70^ analysis of unmapped reads which categorized the greatest number of *Picornaviridae* k-mers to these genomes. Sequences for viral genomes were added to the GRCh38 FASTA and GTF files and processed using Cellranger’s mkref() function. scRNA-seq libraries were then aligned and quantified using Cellranger (v7.2.0) via Cumulus tools^136^. To quantify ambient RNA, Cellbender (v0.2.0)^137^ remove-background() function was applied to each pool with default parameters and epochs = 150, hardware_boot_disk_size_GB = 300, hardware_disk_size_GB = 300, learning_rate = 0.0001, low_count_threshold = 100 and total_droplets_included = 100,000. Using Seurat (v5.1.0)^138^, an object for each pool was created with count matrices for human genes, cellbender corrected human genes, viral genes, and cellbender-corrected viral genes stored in separate assays.

Assignment of RNA+ cells for each virus was done as previously described^15,19^. Briefly, we determined the amount of ambient RNA in each cell by comparing the raw human gene matrices with the cellbender-corrected human gene matrices. Then, we quantified pre-cellbender viral RNA in empty and low-quality droplets (defined as <500 UMI, <300 genes, >35% mitochondrial). Finally, we tested whether the abundance of post-cellbender viral RNA in each cell was significantly above the abundance in empty droplets given the ambient RNA profile of each pool using an exact binomial test (binom.test in R). We performed Benjamini–Hochberg correction to determine the FDR (p.adjust, method = ‘BH’ in R), and cells with FDR < 0.01 were assigned as SARS-CoV-2 RNA+. Following viral RNA+ assignment, non-cellbender human reads were used for all further analysis. Cells were filtered to remove those with <500 UMI, <300 genes, and >35% mitochondrial reads. This resulted in a dataset of 426,577 cells and 40,317 genes across 132 study participants.

#### Single-cell viral metagenomics

Due to challenges identifying Rhinovirus-infected cells in the setting of significant genetic diversity within Enteroviruses and the frequency of coinfections in pediatric patients, we sought to perform an unbiased taxonomic analysis of the data at a single-cell level. Following CellRanger alignment to the human genome, kraken2^70^ was run on unmapped bam files using a previously published kraken2 database that includes all pathogenic human viruses accessible in GenBank as of 2020^139^ using the mg2sc pipeline^140^. The output cell-by-taxon k-mer counts matrix was collapsed to the species level using collapse_taxonomy.py from mg2sc^140^. In R, the .h5ad file was converted to an RDS using Zellkonverter (v.1.10.1)^141^, merged with the GRCh38-aligned Cellranger h5 file, and a Seurat object was created. To create an input file for Cellbender correction, which was used to define true viral-positive cells as above, the Seurat object was saved as an h5 file using DropletUtils (v.1.24.0) Create_10X_h5() function^142^. For downstream analyses, a Seurat object was created with human and kraken reads in separate assays as above. To identify cells containing k-mers classified to specific viruses, species names were identified as follows: for SARS-CoV-2, “Severe acute respiratory syndrome-related coronavirus” and “Severe acute respiratory syndrome coronavirus 2”; for rhinovirus, “Rhinovirus A”, “Rhinovirus B”, and “Rhinovirus C”; for RSV, “Respiratory syncytial virus” and “Human respiratory syncytial virus”. It was noted that a large number of viral k-mers in RSV+ participants were classified at the genus level instead of species level, so an additional classification was created for cells positive for “Human orthopneumovirus” and “Respiratory syncytial virus”. Positive versus negative cells were classified by a binomial test comparing viral k-mers in cells to empty droplets identified by Cellbender, as described in more detail in the preceding section.

#### Evaluation of batch effect

To determine if any batch effects in our dataset necessitated applying integration methods, we first performed sample-level gene expression principal component analysis (PCA). Pseudobulk gene expression profiles from each sample were created using the Seurat AggregateExpression() function and DESeq2 (v1.46.0)^143^. Variance stabilizing transformation was applied before the PCA analysis. To evaluate the impact of metadata variables on the overall variance in the dataset, we pre-processed the Seurat object and performed PCA on the 3,000 most variable human genes. For each principal component (PC), we calculated the proportion of variance explained as the variance of that PC divided by the total variance across the top 20 PCs. We then fit a linear model between each PC and each metadata variable, recording the coefficient of determination (R²). Finally, we estimated the variance contribution of each PC–metadata pair as the product of the PC’s explained variance and the corresponding R². The maximum contribution was between PC1 and participant, explaining 5.1% of the dataset variance **(Supplementary Figure 1F**). Because we did not want to obscure important biological variability between participants, and because we did not observe any significant batch effects based on technical and biological variables of interest, we chose not to integrate our data going forward.

#### Cell clustering and annotation

Non-cellbender-corrected matrices containing only human genes were used in cell annotation. Data were normalized using sctransform (v0.4.1)^144^ with default parameters and method = ‘glmGamPoi’. Cells were clustered using Louvain clustering. Unique marker genes for each cluster were identified using Seurat’s FindAllMarkers() function with cutoffs min.pct = 0.25 and logfc.threshold = 0.25. To assign predicted doublet likelihood scores to each cell, we applied scrublet (v0.2.3)^145^ (with python v3.7.12, scanpy (v1.9.3)^146,147^, anndata (v0.8.0)^147^) to each pool.

We applied an iterative clustering process to remove low-quality cells and confidently determine cell annotations **(Supplementary Figures 2a, 3a**). We first assigned preliminary clusters as either epithelial (n=294,285) or immune (132,292). In each following round, cells of interest were separated and sctransform normalization, PCA, and clustering were repeated. Clusters were removed if 1) the only marker genes were mitochondrial and the percentage of mitochondrial genes was significantly higher than in other clusters, 2) more than 75% of the cells in that cluster came from one pool, indicating a technical artifact, or 3) the cluster was suspected to be doublets based on the expression of marker genes for multiple cell types and higher predicted doublet scores than other clusters. Once objects were divided into cell-type specific objects, clusters were re-assigned if they expressed the marker genes for a different cell type to a greater extent than the marker genes for their original cell type annotation. This approach resulted in the removal of 53,331 epithelial cells (18%) and 40,016 immune cells (30%).

Major epithelial cell types were annotated using marker genes from previously published studies of the human airway^15,19,49–52^. Initial clustering of epithelial cells revealed small clusters of neuronal cells (*RBFOX1*, *CNTNAP2*, *PTPRD*)^148–150^ and sustentacular cells (*ERMN*, *GPX6*, *SEC14L3*)^65^. We next clustered non-ciliated epithelial cells including basal cells (*KRT5, KRT15*), club cells (*SCGB1A1, SCGB3A1*), goblet cells (*MUC5AC, VMO1*), and squamous cells (*SPRR3, SPRR2E*). We identified 5 basal subsets defined by growth genes (*EGFR, SEMA5A*), classical basal markers (*KRT5, TP63*), proliferation genes (*MKI67, TOP2A*), stress-response genes (*EGR1, ATF3*), and secretory cell-like genes (*S100P, KRT7*). There were two club cell subsets alternately expressing interferon (IFN) response genes (*IFIT1*, *IFIT2*) and *BPIFA1*. Exploration of goblet cells revealed 7 subsets: a low-complexity subset we labeled “Developing”, a mitochondrial-high subset, and 5 subsets marked by secretory / mucin genes (*BPIFA1, MUC5B)*, alarmin genes (*S100A8, S100A9*), IFN response genes, statherin (*STATH*), and the cysteine protease inhibitor *CST1*. Squamous cells similarly contained one low-complexity subset, as well as one high for *KRT13* and *HOPX* which resembled “hillock” cells^55,61^ and a pro-inflammatory subset (*IL6, IL1A, CXCL8*).

We identified 7 subsets within ciliated cells (*CAPS, DNAH12)*. 2 ciliated subsets were defined by canonical ciliated markers or cellular growth genes (*MORN5, FOXJ1, GRM7, ESRRG*). The remaining ciliated subsets were defined by class-II antigen presentation genes (*HLA-DPB1*, *HLA-DQA1*), oxidative stress-response genes (*GSTA1, GSTA2*), IFN-response genes (*RSAD2, CXCL11*), keratins and tight junction genes (*KRT17, LAMB3)*, and NF-kB response genes (*NFKBIA, TNFAIP3*). Deuterosomal cells^53,54^ (*CCNO, CDC20B*) contained two ciliated-like subsets (*CROCC, DLEC1, PIFO, MORN2*) and one subset expressing the goblet gene *MUC5AC*. We also identified a population of ionocytes^55,56^ (*CFTR, SCNN1B*) which contained an IFN-responsive subset (*GBP1, CXCL10)*.

We next repeated this iterative clustering with the immune cells. After removing a small cluster of melanocytes (*PMEL*, *MLANA*)^151^, we separated the cells into macrophages (*TYROBP, FCER1G*), plasmacytoid dendritic cells (pDCs) (*IRF7, IRF8, GZMB*), B cells (*MS4A1, BANK1*), mast cells (*CPA3, MS4A2*), and T/NK cells (*CD3D, TRAC, GNLY*). We identified 6 macrophage subsets marked respectively by complement genes (*C1QA, C1QB*), lipid metabolism genes (*APOE, APOC1*), alarmins and chemokines (*S100A8, CCL2*), pro-inflammatory genes (*CXCL8, IL1B*), the signaling molecule *PLCG2*, and a *SPP1*-high subset. We also identified subsets resembling monocyte-derived dendritic cells (*GPR183, CD86*) and conventional type 1 dendritic cells (*CLEC9A*). Plasmacytoid dendritic cells were divided into subsets expressing the transcriptional regulator *ZFAT*, the antimicrobial gene *LYZ* and the receptor tyrosine kinase *AXL*, and high amounts of type I IFNs (*IFNA, IFNB*). B cell subsets included one highly expressing canonical markers *MARCH1* and *PAX5*, a quiescent subset highest in ribosomal genes, an activated subset expressing *NR4A2* and *NR4A3*, an immunoglobulin lambda (*IGLC2, IGLC3*), a cycling subset (*MKI67, TOP2A*), and a distinct subset of plasma cells (*MZB1, JCHAIN, IGHA1*). We next separated natural killer (NK) cells from T cells based on their phenotype and lack of *CD3D* expression. We identified 3 NK cell subsets: a quiescent ribosomal-high subset, an *NCAM1*+ high subset, and a *CCL4*+ subset. We also noted a small cluster of innate lymphoid cells resembling lymphoid tissue inducer (LTi) cells based on the expression of *IL4I1, KIT, IL23R,* and *LTB*. We then defined gamma-delta (γδ) T cells as those that expressed both *TRGC2* and *TRDC* more highly than *TRAC* and *TRBC2*. There were 4 subsets of γδ T cells expressing ribosomal genes, the transcriptional repressor *ZBTB20*, immune signaling molecules *XCL1* and *XCL2*, and IFN-responsive genes respectively. When we sub-clustered alpha-beta (ab) T cells, we found that many phenotypic clusters (such as resident-memory T (Trm) cell-like, cycling, quiescent, and cytotoxic) contained both CD4+ and CD8+ T cells. To elucidate both the T cell lineage and the phenotype, we first separated the ab T cells into 7 phenotypic clusters. Next, we sub-clustered each of these to divide each one into CD4+ and CD8+ cells. Finally, we re-combined all identified CD4+ and CD8+ cells, respectively, and performed the final sub-clustering and annotation. Within CD4+ T cells, we found a quiescent subset, a naive-like *IL7R+ KLRB1+* subset, a subset expressing exhaustion-related genes (*HAVCR2, CD38*), an activated *TNFRSF4+* subset, Th17 cells (*IL17A, IL23R*), Tph cells highly expressing *CXCL13*, and Tregs (*FOXP3, IL2RA*). With CD8+ T cells, we similarly find quiescent and ribosomal-high subsets, a Trm-like subset (*ITGAE, ITGA1*), an IFN-responsive subset, chemokine-high *CCL4+IFNG+* cells, conventional cytotoxic T cells (*GZMB, PRF1*) as well as a subset highly expressing granulysin (*GNLY*). In addition, we found two small subsets of CD8+ T cells expressing *IL17A* and *CXCL13*, respectively. Finally, we identified a cluster of MAIT cells (*ZBTB16*, *SLC4A10*) as well as cycling CD4+ and CD8+ T cells. Marker genes used to annotate clusters in the final round for each cell type are provided in **Supplementary Table 3**.

#### Pseudobulk differential expression

To compare transcriptional signatures between different groups of participants or cells within a subset of interest, we created pseudobulk aggregates for each participant using Seurat’s AggregateExpression(). A DESeqDataset (DESeq2 v.46.0) object was then constructed from the aggregated count matrix, and genes with <10 counts across all samples were filtered out. Differential expression testing was done using the DESeq() function with default parameters and contrasts specified per analysis. Genes with an adjusted p-value (FDR) <0.05 were considered statistically significant. RSV+ CXCL11+ IFN-stim ciliated cells vs. SARS-CoV-2+ CXCL11+ IFN-stim ciliated cells **(Figure 2F**): participants with at least 20 CXCL11+IFN-stim cells were included (SARS-CoV-2: n=4; RSV: n=6); RSV+ KRT17+S100A9+ ciliated cells vs. RSV-KRT+S100A9+ ciliated cells **(Figure 3B**): participants with at least 20 KRT17+S100A9+ ciliated cells were included (n=27); T cells in ICU admitted RSV participants vs. T cells in floor-admitted RSV participants **(Figure 6E**): all admitted RSV participants were included (admitted to floor: n = 14, admitted to ICU: n =7). Complete lists of DE genes for each of these comparisons are provided in **Supplementary Table 4**.

#### RNA velocity

We compared potential infected cell fates using RNA velocity with scVelo (v0.3.3)^152^. Briefly, loom files containing unspliced and spliced RNA matrices were created by running velocyto (v0.17.17)^153^ using Cellranger-aligned bam files, the GRCh38-2020-A reference genome (Ensembl 93), and GRCh38 with repetitive elements masked. The masked GRCh38 genome was obtained from the UCSC genome browser (https://genome.ucsc.edu/cgi-bin/hgTables) by downloading the gtf file of GRCh38 using track “Repeat Masker” for “All tracks”^154^. Aggregated loom files were merged with the processed, clustered, and annotated epithelial Anndata object and subset to high-quality cells from Control, SARS-CoV-2, and RSV groups. Following scVelo filtering, normalization, nearest neighbor calculation, and moments computation, splicing dynamics were calculated using the dynamical model. The velocity streams were visualized in UMAP space.

#### Transcription factor scores

Transcription factor scores were calculated by univariate linear modeling (ULM) of SCT-transformed counts with the collecTRI database^155^ of transcription factor regulons using the decoupler package(v2.12)^156^ in Python. Enriched transcription factor activities for subsets of interest compared to all other subsets were selected by t-test-overstim-var within decoupler.

#### Compositional analysis

For compositional analyses, we first created an abundance matrix of either cell types or cell subsets for each participant. Center-log-ratio (CLR) normalization was carried out using the setaCLR() function from the SETA (v1.0.0)^157^ package. To assess changes in cell type abundance relative to controls, we calculated the difference between the average CLR-normalized abundance for each viral group compared to the average CLR-normalized abundance for the control group. PCA on the CLR-normalized cell subset abundance matrix was computed using the setaLatent() function, method = “PCA”, dims=10. To evaluate relationships between cell subsets, Spearman correlation between the CLR-normalized abundance for each combination of cell subsets was calculated.

#### Cytokine response signature scores

Response signatures were derived from gene expression of cytokine-treated airway basal cells^72,73^ (**Figure 3E-F**, **Figure 4D-E, Supplementary Figure 5C–D**; IFNα, IL-4, and IL13 response signatures in epithelial cells), from the Hallmark database using msigdbr (v25.1.1)^158^ **(Figure 6F, Supplementary Figure 8F**; IFNα and IFNγ response signatures in all cells), or gene response signatures from glucocorticoid treatment of airway epithelial cells^92^ **(Supplementary Figure 8G**). Gene signature scores per participant were calculated using univariate linear modeling (ULM) of pseudobulk-averaged SCT-transformed counts using the R implementation of decoupler^156^ (v2.9.7). To score gene signatures on a per-cell level **(Supplementary Figure 5C**), univariate linear modeling was applied to a downsampled object containing 10% of cells from each epithelial cell subset. Genes included in each signature are provided in **Supplementary Table 5**.

#### Cell-cell communication

To assess differences in cell-cell communication between groups, we applied multinichenetr (v2.1.0)^89^, an extension of nichenetr (v2.2.1)^159^ designed for multi-sample, multi-condition differential analysis. Pseudobulk expression was calculated for each sample and cell type combination. The contrast design compared the control group to all other samples and each viral group to controls only. We did not define any batch or covariate variables. When evaluating the differential expression of ligands of interest **(Figure 5B**), we used more permissive cutoffs for the minimum number of cells per sample-cell type combination(min_cells = 5) and the minimum proportion of samples in which a gene was expressed (min_sample_prop = 0.25) and recommended cutoff for fraction of cells with nonzero expression per sample-cell type combination (fraction_cutoff = 0.05). Out of a comprehensive list of cytokines and chemokines **(Supplementary Table 6**), we evaluated which were identified as differentially expressed (FDR<0.05) in each viral group compared to controls.

To prioritize receptor-ligand interactions, we applied standard cutoffs to filter cell types and genes included in the pseudobulk (min_cells =10, sample_prop = 0.5, fraction_cutoff = 0.05). The same contrast design described above was used to identify differentially expressed genes for each cell type. To assess ligand activities, we considered the top 250 target genes and standard parameters. All prioritization criteria were weighed equally. The top 50 ranked interactions were plotted in **Figure 5C** and **Supplementary Figure 6A**, though we evaluated the number of interactions, the prioritization scores, and the distribution of sender cell types for various cutoffs **(Supplementary Figure 6B-D**). To visualize prioritization metrics, we calculated the average scaled pseudobulk expression product of the ligand and receptor for each group and interaction. We narrowed the interactions displayed in **Figure 5D** by 1) selecting the sender cell types that accounted for at least 5% of interactions and 2) showing only the highest ranked interaction for repeated receptor-ligand-receiver combinations. The complete list of the top 500 ranked interactions can be found in **Supplementary Table 6**.

#### Predictive modeling of cell subset importance

We trained a penalized regression classifier with 5-fold cross-validation to distinguish patients by Wheeze_Group based on cell subset abundance using the glmnet model from the Caret package (v7.0.1)^160^. Variables (cell subsets) that contributed the most to predicting Wheeze_Group were extracted for plotting.

#### scDRS analysis

We linked our single cell transcriptomes with polygenic disease risk for asthma and height using scDRS (v1.0.4)^97^. GWAS summary statistics for childhood-onset asthma (GCST007800) and adult-onset asthma (GCST007799)^161^ were downloaded from the NHGRI-EBI GWAS Catalog^162^ on 2025-09-16. SNP-level summary statistics were converted to gene-level Z-scores using MAGMA version (v1.1.0)^163^. We removed HLA regions (chr6: 25-34 Mb) and used the 1000 Genomes European cohort as our reference panel. Gene-level scores were converted to scDRS .gs format with scDRS munge-gs (--weight zscore, ––n-max 1000, and ––n-min 100) and merged with pre-processed gene sets provided by the original scDRS study.

We then ran scDRS using a Nextflow workflow set up for this analysis using default parameters. During preprocessing, we corrected for Batch, Collection_Site, and LeviCell using scdrs.pp.preprocess (n_mean_bin=20, n_var_bin=20). Cell-level scores were obtained with scdrs.method.score_cell (n_ctrl=1000 and ctrl_match_key=“mean_var”). For downstream analysis, we used scdrs.method.downstream_group_analysis, scdrs.method.downstream_corr_analysis, and scdrs.method.downstream_gene_analysis.

#### Label transfer

Data was aligned using BeeNetPlus (v1.0.X) software implemented on Terra (Honeycomb Biotechnologies). Seurat objects for each independent cohort were created from raw count matrices (tcm.tsv files) for each sample. For the BD-rural cohort, samples were aligned to a custom genome including both human and RSV genes designed for BeeNetPlus software. Count matrices for viral genes were stored in a separate assay within the Seurat object. Cells with <100 transcripts, <100 unique genes, or >35% mitochondrial reads were filtered out. Objects were first clustered independently at low resolution to assign cells as epithelial or immune. Each epithelial / immune object was then clustered once to ensure the correct identity and to remove any low-quality cells. This resulted in high-quality epithelial (IND-Kolkata cohort: n=16,523 cells; BD-rural cohort: n=12,715 cells) and immune (IND-Kolkata cohort: n=40,544 cells; BD-rural cohort: n=5,266 cells) objects.

Labels were then transferred within each lineage. For the IND-Kolkata cohort, only samples in the control group in the USA-Boston cohort were used as a reference. For the BD-rural cohort, labels were transferred from all samples in the USA-Boston cohort. UMAP embedding for the reference object was first re-calculated based on the cells and samples included. Label transfer anchors were identified using the FindTransferAnchors() function with the union of variable genes as the features of interest and normalization.method = “SCT”, then used as the anchorset in the TransferData() function. The anchorset was also used to project the UMAP embedding from the reference object to the query object via MapQuery(). Label transfer was evaluated by examining the distribution of prediction scores and consistency with manual annotations. Similarity in abundance of cell subsets between reference and query datasets was evaluated by 1) calculating the frequency of each subset out of all epithelial or immune cells per sample, 2) taking the average frequency within all samples from one site, 3) ranking the subsets by their average frequency such that the most abundant subset had the highest ranking and 4) correlating the rankings between each site.

## Supplementary Figures

**Supplementary Figure 1.**
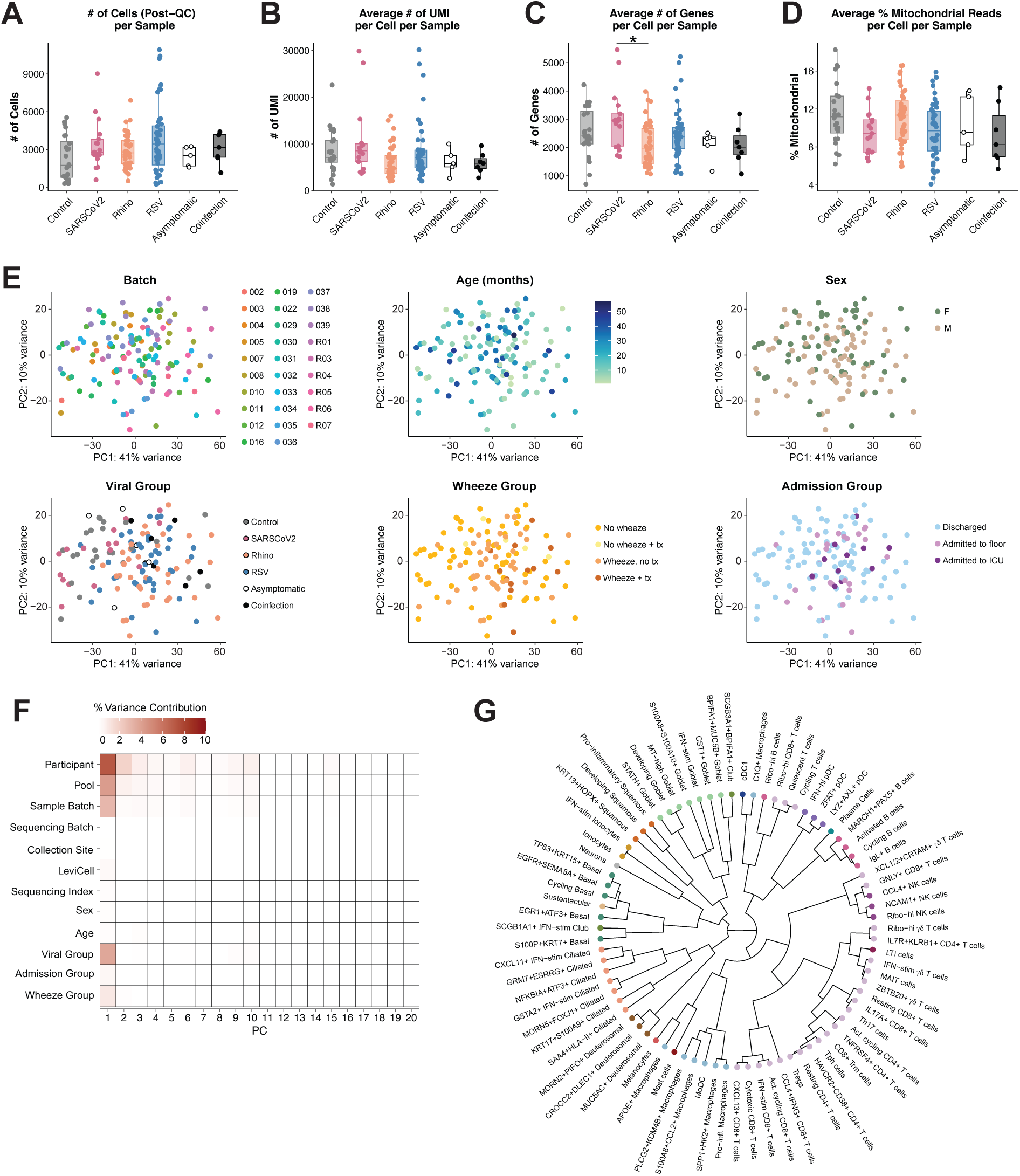
Related to Figure 1. **A.** Number of cells per sample after selecting high-quality cells (defined as >500 unique molecular identifiers (UMI), >300 unique genes, and <35% mitochondrial reads) by viral group. Metrics in **B-D** are restricted to high-quality cells. QC = quality control; Rhino = rhinovirus. **B.** Average number of UMI per cell per sample by viral group. **C.** Average number of unique genes per cell per sample by viral group. **D.** Average percentage of mitochondrial reads per cell per sample by viral group. **E.** Principal component analysis (PCA) of pseudobulk aggregated gene expression for each sample, colored by metadata variables. F = female; M = male; tx = wheezing-directed medication; ICU = intensive care unit. **F.** Variance contribution matrix for metadata variables of interest and top 20 principal components (PC) from single-cell gene expression data. Color represents the percentage of variance in the dataset explained by each variable through each PC. **G.** Hierarchical tree of cell subsets as in **Fig. 1I** with subset labels. End nodes are colored by cell type of origin. Hierarchical distances were calculated from pseudobulk aggregates of each cell subset. IFN-stim = interferon-stimulated; MT = mitochondrial; MoDC = monocyte-derived dendritic cell; cDC1 = conventional type 1 dendritic cell; pDC = plasmacytoid dendritic cell; Ribo. = ribosomal; LTi = Lymphoid tissue inducer cell; MAIT = mucosal-associated invariant T cell; Th17 = T helper cell type 17; Act. = activated; Trm = resident memory T cell; Tph = T peripheral helper cell; Treg = regulatory T cell. **A-D:** Boxplots represent the median (center line), interquartile range (box), and 1.5x the interquartile range (whiskers). Statistical test is Kruskal-Wallis test with Benjamini-Hochberg correction for multiple comparisons. Dunn’s post-hoc test, *p<0.05

**Supplementary Figure 2.**
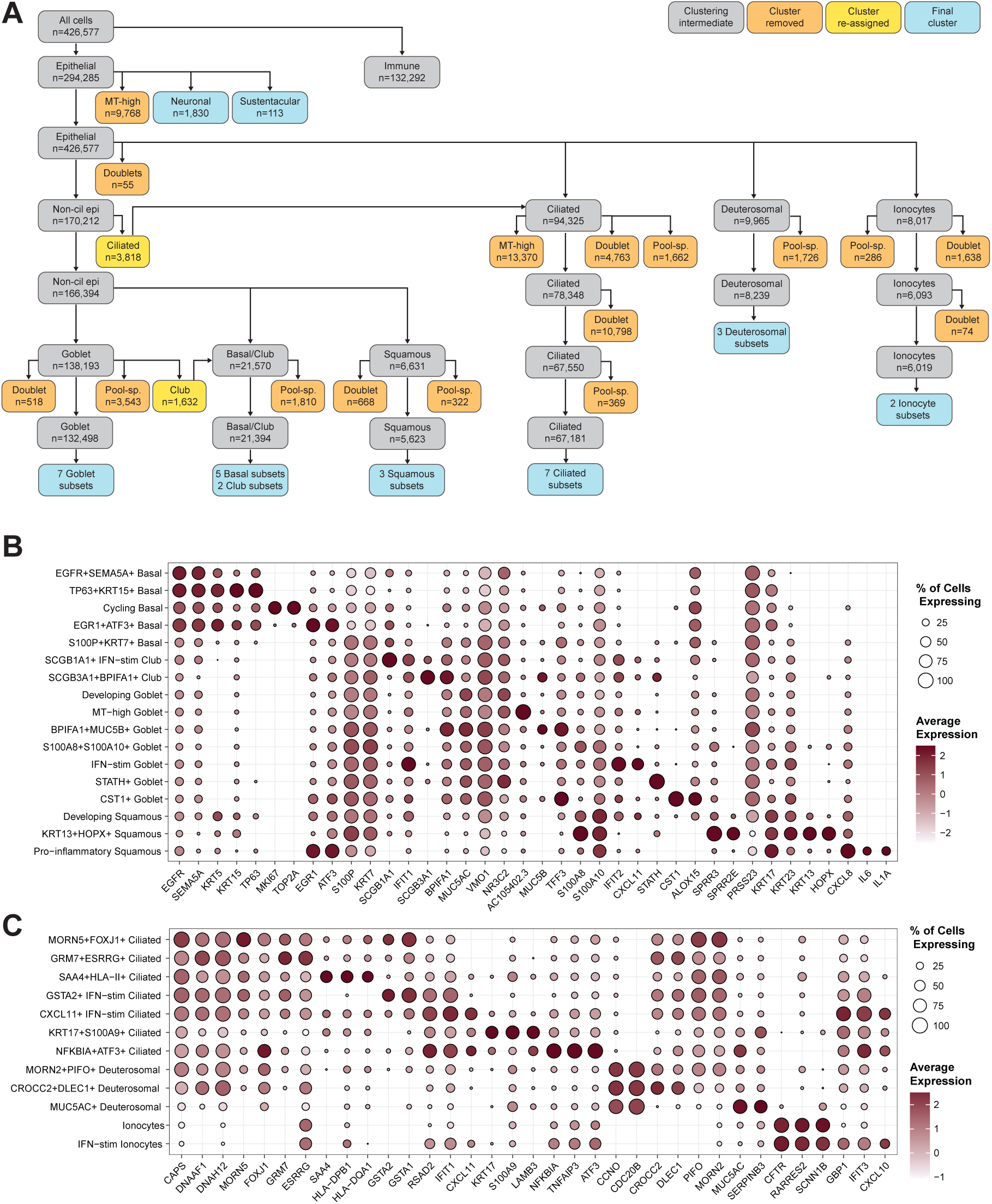
Related to Figure 1. **A.** Schematic of epithelial cell clustering approach. Clusters were removed at each level if they were defined by only mitochondrial genes, had >75% of cells from one sample pool, or were suspected doublets (see **Methods**). Clusters were re-assigned if they expressed marker genes for a different cell type to a higher degree than the marker genes for the cell type of interest. Normalization was repeated at each level. Final clusters were determined after comparing multiple resolutions. Pool-sp = pool-specific, Non-cil = non-ciliated. **B.** Marker genes for basal, club, goblet, and squamous cell subsets. IFN-stim = interferon-stimulated. **C.** Marker genes for ciliated, deuterosomal, and ionocyte subsets. **B-C.** See **Supplementary Table 3** for complete lists of marker genes for final cell subsets.

**Supplementary Figure 3.**
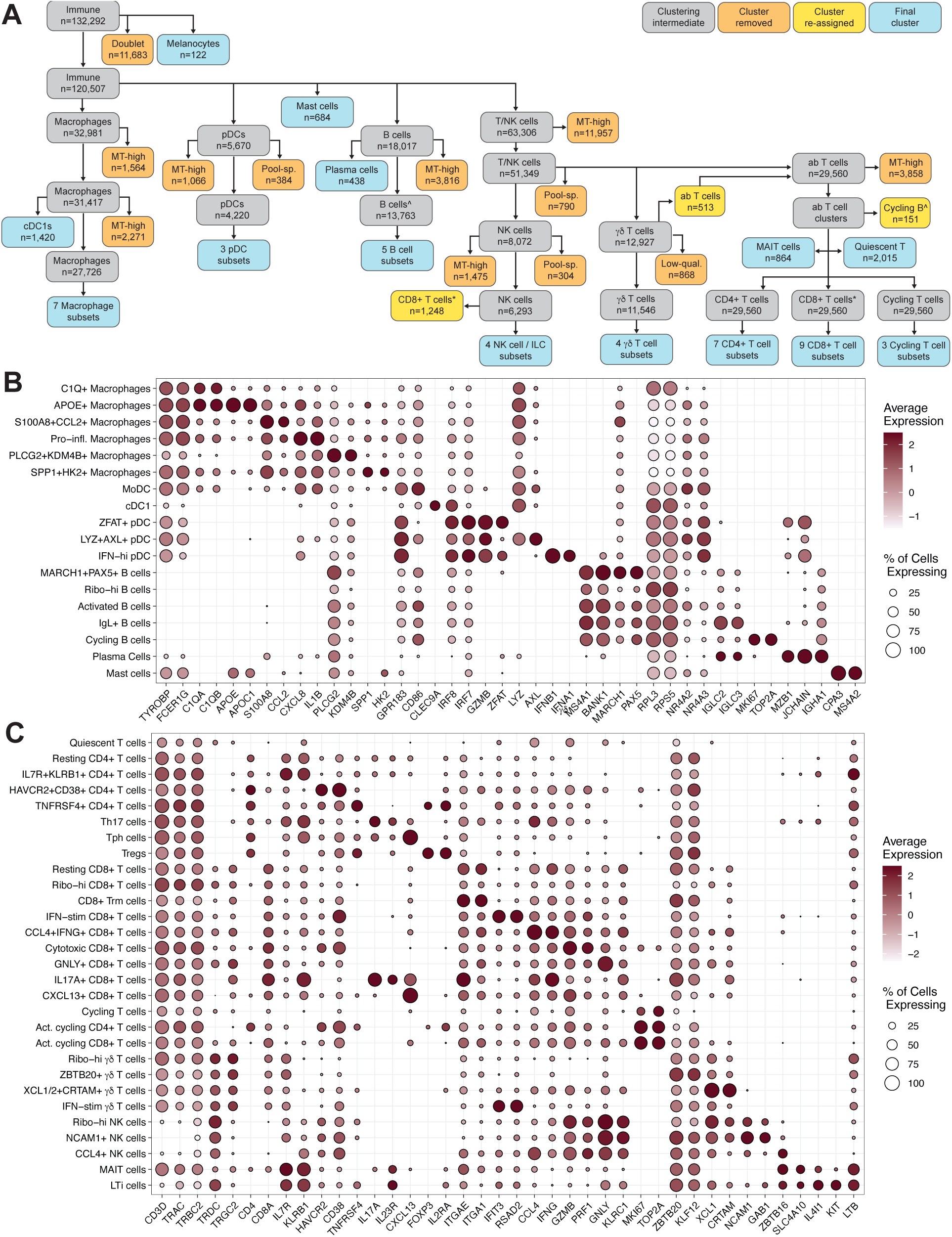
Related to Figure 1. **A.** Schematic for immune cell clustering approach. See legend for **Supplementary Figure 2** for further information. Pool-sp = pool-specific.^* denotes where re-assigned clusters were included if not shown with arrows. **B.** Marker genes for macrophage, dendritic cell, B cell, and mast cell subsets. MoDC = monocyte-derived dendritic cell; cDC1 = conventional type 1 dendritic cell; pDC = plasmacytoid dendritic cell. **C.** Marker genes for T cell and natural killer (NK) cell subsets. Th17 = T helper cell type 17; Tph = T peripheral helper cell; Treg = regulatory T cell; Ribo. = ribosomal; Trm = resident memory T cell; Act. = activated; IFN-stim = interferon-stimulated; MAIT = mucosal-associated invariant T cell; LTi = Lymphoid tissue inducer cell. **B-C.** See **Supplementary Table 3** for complete lists of marker genes for final cell subsets.

**Supplementary Figure 4.**
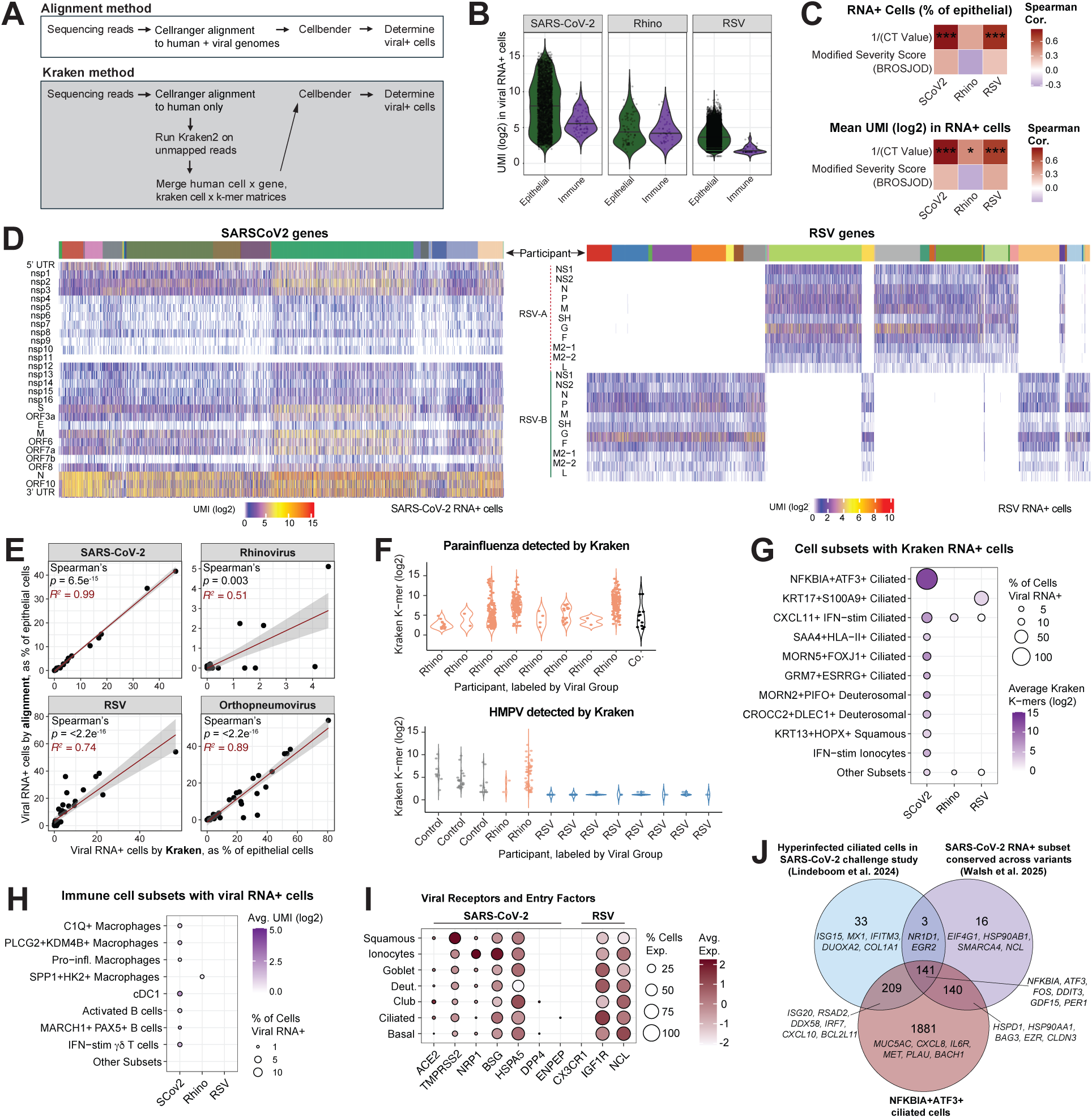
Related to Figure 2. **A.** Schematic for identifying viral RNA+ cells by alignment or Kraken2 metagenomics approach. See Methods for details. **B.** Violin plot demonstrating viral UMI/cell for epithelial compared to immune viral RNA+ cells for SARS-CoV-2, rhinovirus, and RSV. Rhino = rhinovirus. RSV = respiratory syncytial virus. **C.** Spearman correlation between viral RNA+ cells as a percentage of epithelial cells and inverse CT value from qRT-PCR of supernatant collected during swab processing and disease severity (see **Methods** for details) for each virus (top) and modified BROSJOD disease severity score (see **Methods** for details, bottom). SCoV2 = SARS-CoV-2. Spearman correlation test with Benjamini-Hochberg correction for multiple comparisons. *p<0.05, ***p<0.001. **D.** Heatmap of SARS-CoV-2 (left) and RSV (right) gene expression in viral RNA+ cells. Top bar depicts the individual participant for each cell. UMIs shown are post removal of background reads by Cellbender. **E.** Spearman correlation between viral RNA+ cells as a percentage of epithelial cells between alignment and Kraken methods for respective viral groups. R^2^ and Spearman *p* values are shown. For RSV, two different Kraken2 levels of classification are depicted. The plot labeled “RSV” indicates the custom kraken database species-level identifier “Respiratory Syncytial Virus” compared to RSV+ cells by alignment method. The plot labeled “Orthopneumovirus” indicates the custom kraken database genus-level identifier “Orthopneumovirus” compared to RSV+ cells by alignment method. **F.** Violin plot of kraken k-mers for parainfluenza or human metapneumovirus (HMPV) per participant for participants with >1 viral RNA+ cell for parainfluenza or HMPV. Participants are labeled by viral group. Co. = coinfection. **G.** Dotplot of viral RNA+ cells identified by Kraken method in all subsets for SARS-CoV-2, rhinovirus, and RSV. Dot size represents the percentage of cells in the subset that are viral RNA+. Color represents the average viral k-mers across all cells in the subset. Cell subsets with <5% viral RNA+ cells are collapsed. Cell subsets with fewer than 100 cells/group were excluded. **H.** Dotplot of viral RNA+ cells by alignment method in each immune subset for SARS-CoV-2, rhinovirus, and RSV. Dot size represents the percentage of cells in the subset that are viral RNA+. Color represents the average viral transcripts (UMI) across all cells in the subset. Cell subsets with <1% viral RNA+ cells are collapsed. Cell subsets with fewer than 100 cells/group were excluded. **I.** Dotplot of SARS-CoV-2 and RSV viral receptor and entry factors within each epithelial cell type. Includes all epithelial cells from all participants. **J.** Venn diagram comparing marker genes for *NFKBIA+ATF3+* ciliated cells in this dataset, “hyperinfected” ciliated cells identified in a SARS-CoV-2 human challenge study^18^ and a PER2+EGR1+GDF15+ ciliated subset consistently enriched in viral RNA+ cells across SARS-CoV-2 variants^19^.

**Supplementary Figure 5.**
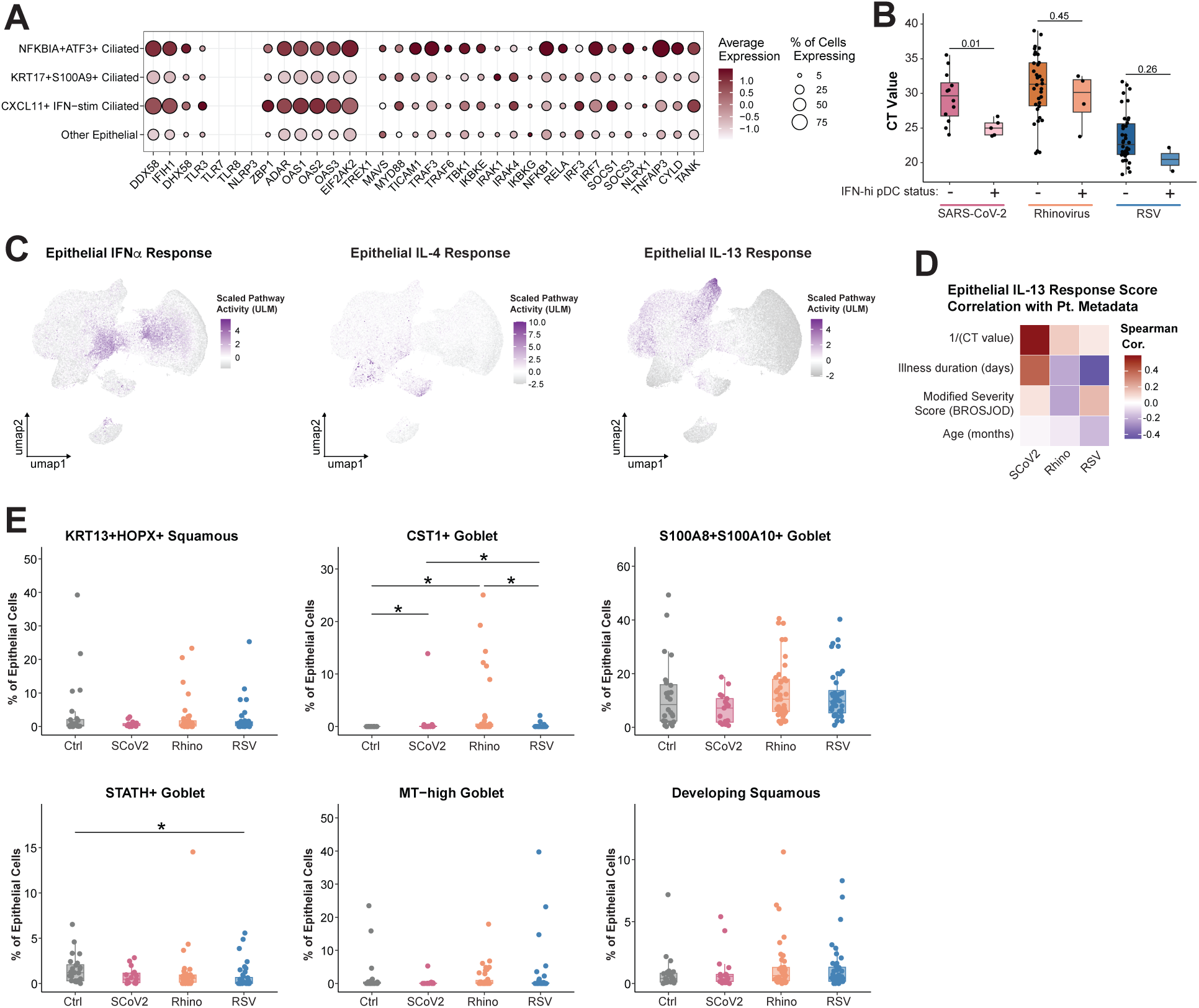
Related to Figures 3 and 4. **A.** Dotplot of pattern recognition receptor and antiviral genes in viral RNA+ epithelial cell subsets. **B.** Viral RNA CT value from supernatant collected during swab processing divided by viral group and IFN-hi plasmacytoid dendritic cell status (+ = >0 IFN-hi pDCs in the sample). Statistical test is a Mann-Whitney U test. **C.** UMAP of all epithelial cells from participants in control and mono-infection groups colored by scaled pathway activity score for IFNα (left), IL-4 (center), and IL-13 (right) response genes. Scoring was done on a downsampled object containing 10% of cells from each epithelial cell subset. Signatures provided in **Supplementary Table 5**. **D.** Spearman correlation between pseudobulk epithelial IL-13 pathway activity score and select participant metadata variables within each viral group. CT value represents viral RNA CT value from supernatant collected during swab processing. Spearman correlation test with Benjamini-Hochberg correction for multiple comparisons. *p<0.05, otherwise n.s. **E.** Frequency of specified epithelial subsets as a percentage of epithelial cells per sample, compared across viral groups. Participants with <100 epithelial cells were excluded. Statistical test is Kruskal-Wallis test with Benjamini-Hochberg correction for multiple comparisons. Dunn’s post-hoc test, *p<0.05. **B, E:** Boxplots represent the median (center line), interquartile range (box), and 1.5x the interquartile range (whiskers).

**Supplementary Figure 6.**
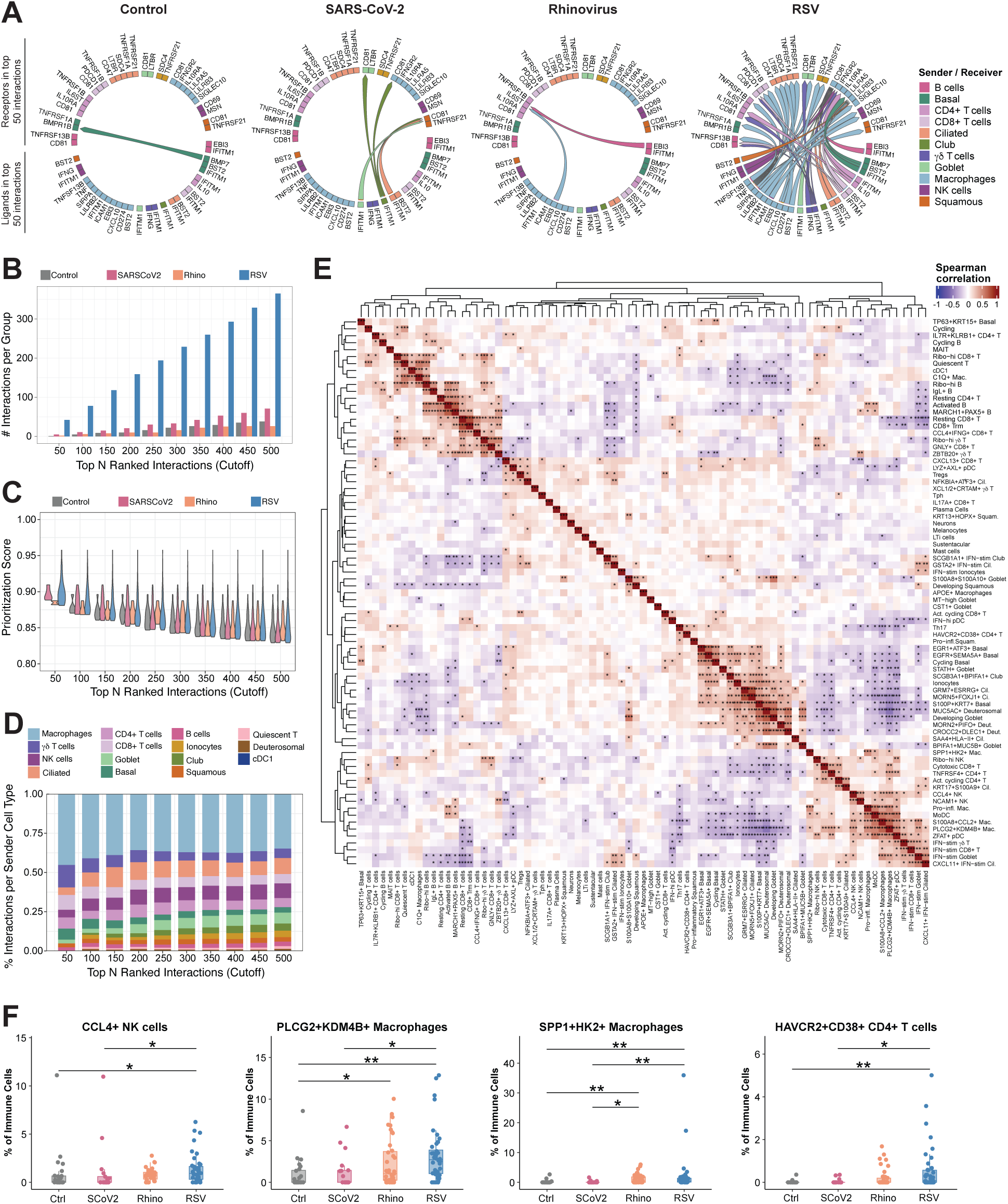
Related to Figure 5. **A.** Chord diagrams of top 50 overall ranked cell-cell interactions (multinichenet) as in **Figure 5C**, including ligand receptor labels. Starting point of chord represents the ligand, endpoint represents the receptor. Chords are colored by sender cell type. **B.** Number of interactions specific to each viral group in top interactions across various cutoffs. See **Supplementary Table 6** for list of top 500 prioritized interactions. **C.** Prioritization scores of interactions specific to each viral group in top interactions across various cutoffs. **D.** Distribution of most frequent sender cell types in top interactions across various cutoffs. **E.** Spearman correlation of CLR-normalized cell subset abundances among all samples as in **5E**, including subset labels. Both rows and columns were clustered using hierarchical clustering with Euclidean distance and complete linkage. P values from correlation test adjusted with Benjamini-Hochberg correction for multiple comparisons. *p<0.05, **p<0.001, ***p<0.0001. **F.** Frequency of specified immune subsets as a percentage of immune cells per sample, compared across viral groups. Participants with <100 immune cells were excluded. Boxplots represent the median (center line), interquartile range (box), and 1.5x the interquartile range (whiskers). Statistical test is Kruskal-Wallis test with Benjamini-Hochberg correction for multiple comparisons. Dunn’s post-hoc test, *p<0.05, **p<0.001.

**Supplementary Figure 7.**
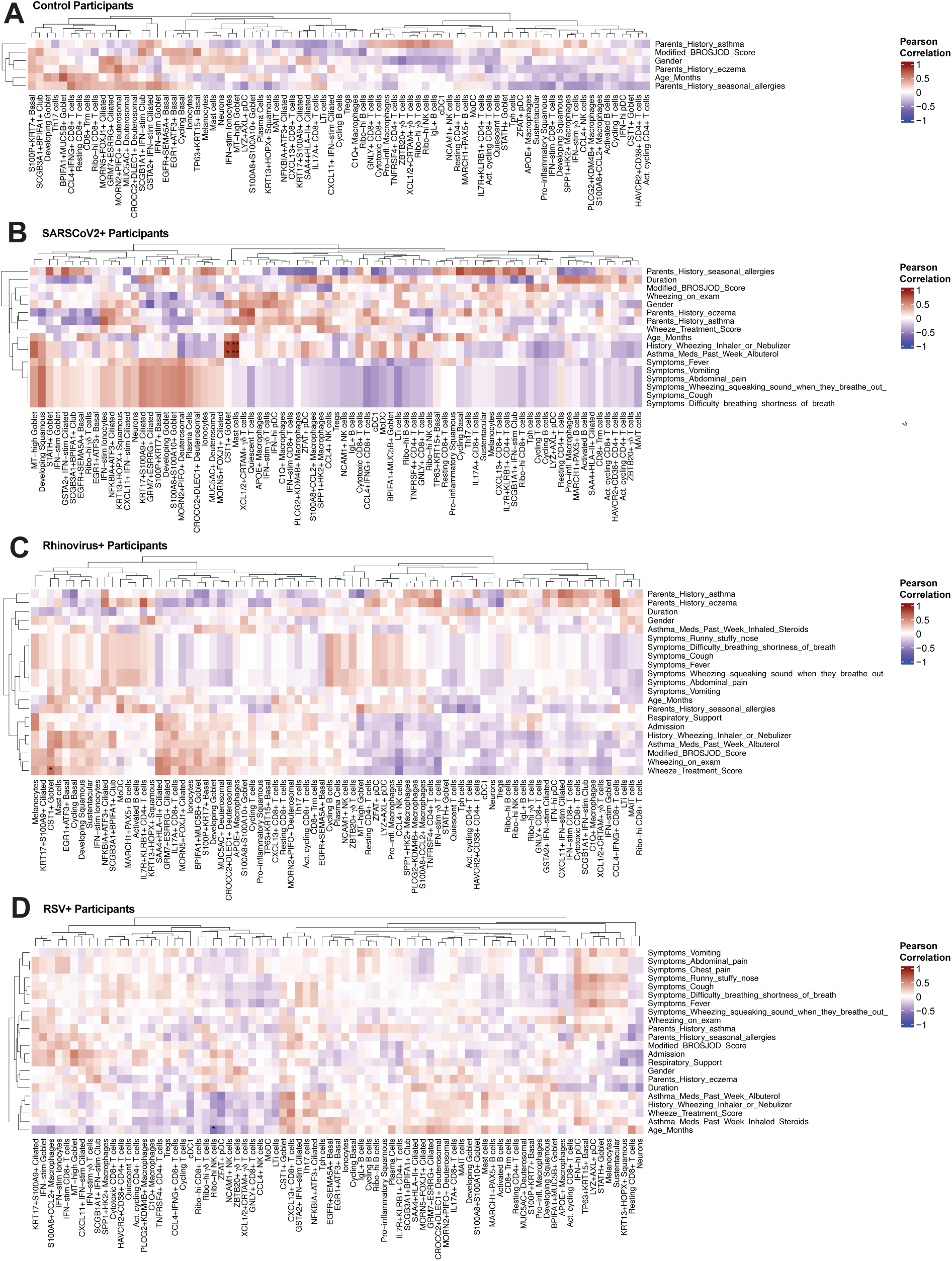
Related to Figure 6. **A-D.** Pearson correlation of CLR-normalized cell subset abundances among **(A**) control, **(B**), SARS-CoV-2, **(C**) Rhinovirus participants, and **(D**) RSV participants. Both rows and columns were clustered using hierarchical clustering with Euclidean distance and complete linkage. Pearson correlation test with Benjamini-Hochberg correction for multiple comparisons. *p < 0.05, **p < 0.01.

**Supplementary Figure 8.**
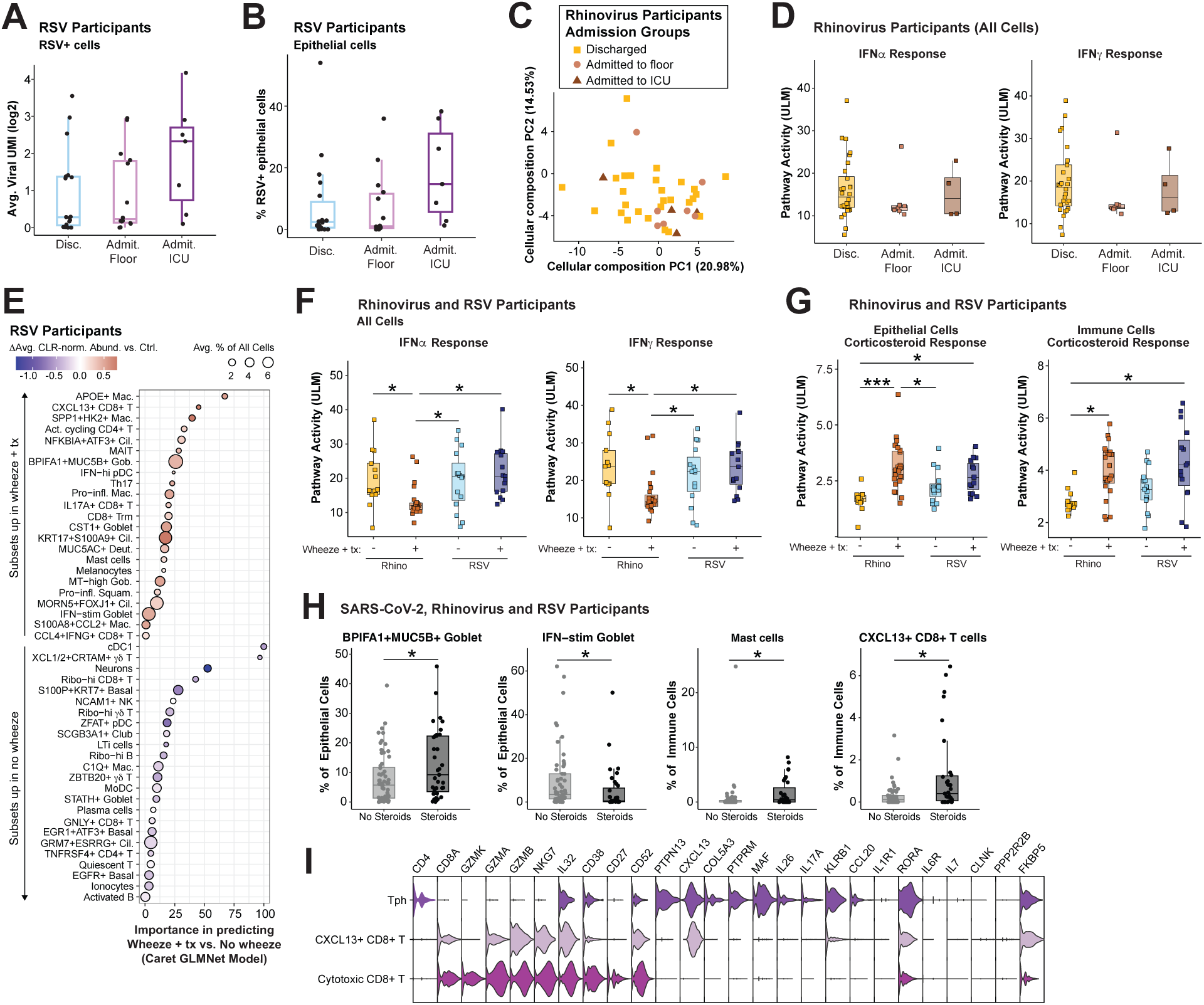
Related to Figure 6. **A.** Average number of viral UMI in RSV+ cells for each RSV participant, compared across admission groups. Disc. = discharged; Admit. Floor = admitted to floor, Admit. ICU = admitted to intensive care unit. **B.** Frequency of RSV+ cells out of all epithelial cells per sample, compared across admission groups. **C.** Principal component analysis (PCA) of cellular composition for each sample in rhinovirus group colored by admission groups. PCA was run on center-log-ratio (CLR) normalized cell subset abundances. See **Figure 3G** for PC loadings. **D.** Pseudobulk pathway activity score for IFNɑ (left) and IFNɣ (right) response genes per Rhinovirus participant in all cells, compared across admission groups. Signatures provided in **Supplementary Table 5**. **E.** Machine-learning-derived subset importance for distinguishing no wheeze vs wheeze + treatment for RSV participants by Caret’s penalized generalized linear (GLMNet) model. All subsets that contributed to model importance are shown. Color of dot indicates the difference in center-log-ratio (CLR)-normalized cell subset abundance between patients with wheeze+tx compared to no wheeze. Dot size represents frequency of the cell subset, averaged across all included RSV participants. Tx = treatment. Mac. = macrophage; Cil. = ciliated; MAIT = mucosal-associated invariant T; Gob. = goblet; pDC = plasmacytoid dendritic cell; Th17 = T-helper 17; Trm = T resident memory; Deut. = deuterosomal; Squam. = squamous; cDC = conventional dendritic cell; NK = natural killer; LTi = lymphoid tissue inducer; MoDC = monocyte-derived dendritic cells. **F.** Pseudobulk pathway activity score for IFNɑ (left) and IFNɣ (right) response genes per participant in Rhinovirus and RSV groups in all cells, compared across wheeze groups (no wheeze/no treatment (-) or wheeze + treatment (+)). Tx = treatment. **G.** Pseudobulk pathway activity score for corticosteroid response genes per-participant score in Rhinovirus and RSV groups in epithelial (left) and immune cells (right), compared across wheeze groups (no wheeze/no treatment (-) or wheeze + treatment (+)). Signature provided in **Supplementary Table 5**. **H.** Frequency of specified subsets as a percentage of epithelial or immune cells for all participants in SARS-CoV-2, Rhinovirus, and RSV groups, compared by treatment with corticosteroids in the week prior to presentation or in the ED. **I.** Violin plot of marker genes for *CXCL13+* CD8 T cells, T peripheral helper (Tph) cells, and cytotoxic CD8 T cells. **A,B,D,F,G,H:** Boxplots represent the median (center line), interquartile range (box), and 1.5x the interquartile range (whiskers). **A,B,D,F,G,H:** Statistical test is Kruskal-Wallis test with Benjamini-Hochberg correction for multiple comparisons. Dunn’s post-hoc test, *p<0.05, **p<0.001, ***p<0.0001, otherwise n.s.

**Supplementary Figure 9.**
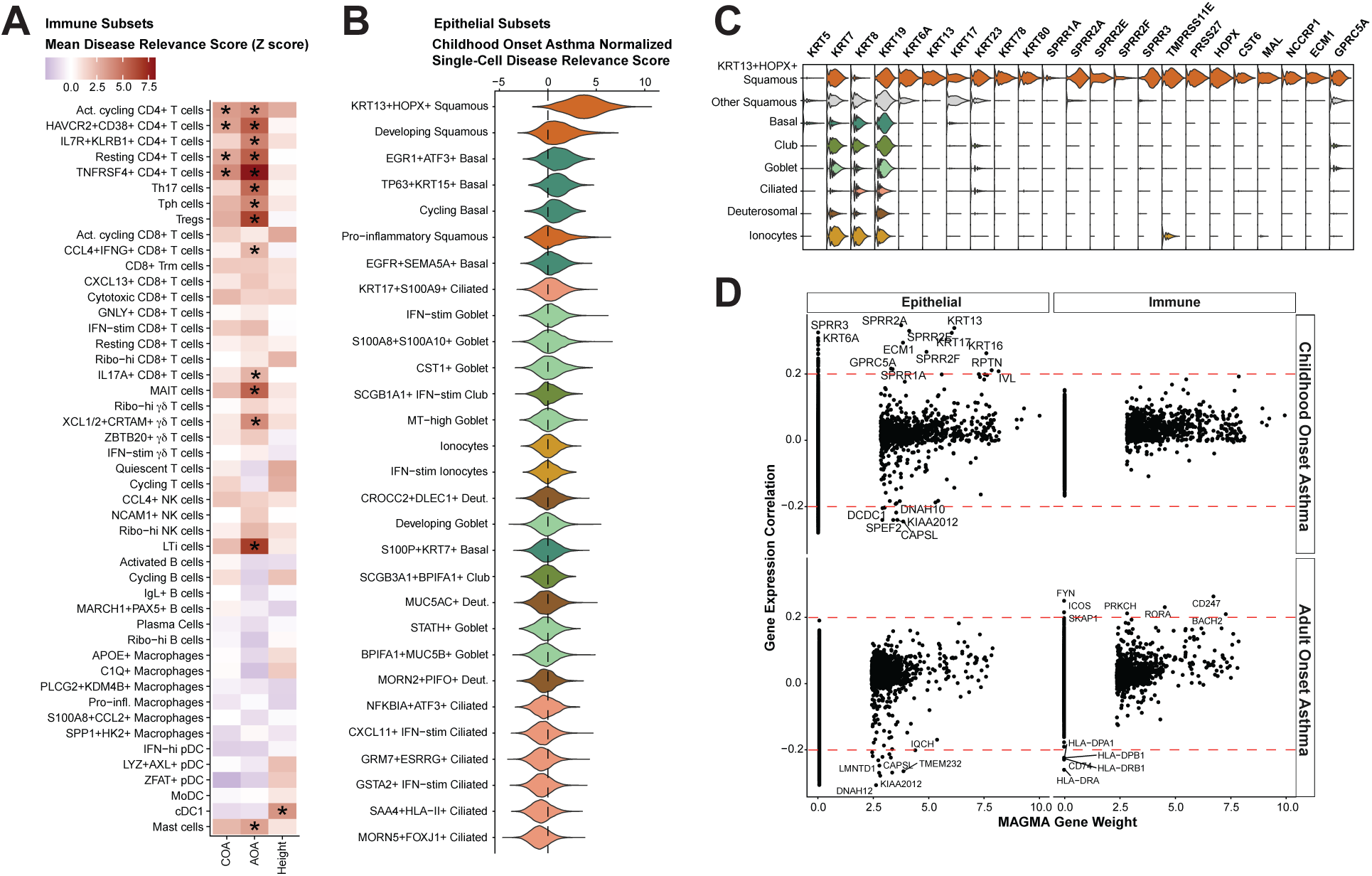
Related to Figure 7. **A.** Cell type-trait association Z-score for childhood onset asthma (COA), adult onset asthma (AOA), and height across immune subsets. *FDR adjusted p<0.05. NK = natural killer cell. Th17 = T helper cell type 17; Tph = T peripheral helper cell; Treg = regulatory T cell; Ribo. = ribosomal; Trm = resident memory T cell; Act. = activated; IFN-stim = interferon-stimulated; MAIT = mucosal-associated invariant T cell; LTi = Lymphoid tissue inducer cell, MoDC = monocyte-derived dendritic cell; cDC1 = conventional type 1 dendritic cell; pDC = plasmacytoid dendritic cell. **B.** Violin plot of scDRS z-scores for childhood onset asthma by epithelial cell subset. Colors represent epithelial cell types. Deut. = deuterosomal. **C.** Marker genes for *KRT13+HOPX+* squamous epithelial subset. **D.** Relationship between gene expression-trait correlation and MAGMA gene weights for that trait, split by epithelial (left) and immune (right), childhood onset asthma (top) and adult onset asthma (bottom). Dotted lines mark absolute correlations greater than 0.2. Genes with a 0 MAGMA gene weight had no GWAS association with the disease trait.

**Supplementary Figure 10.**
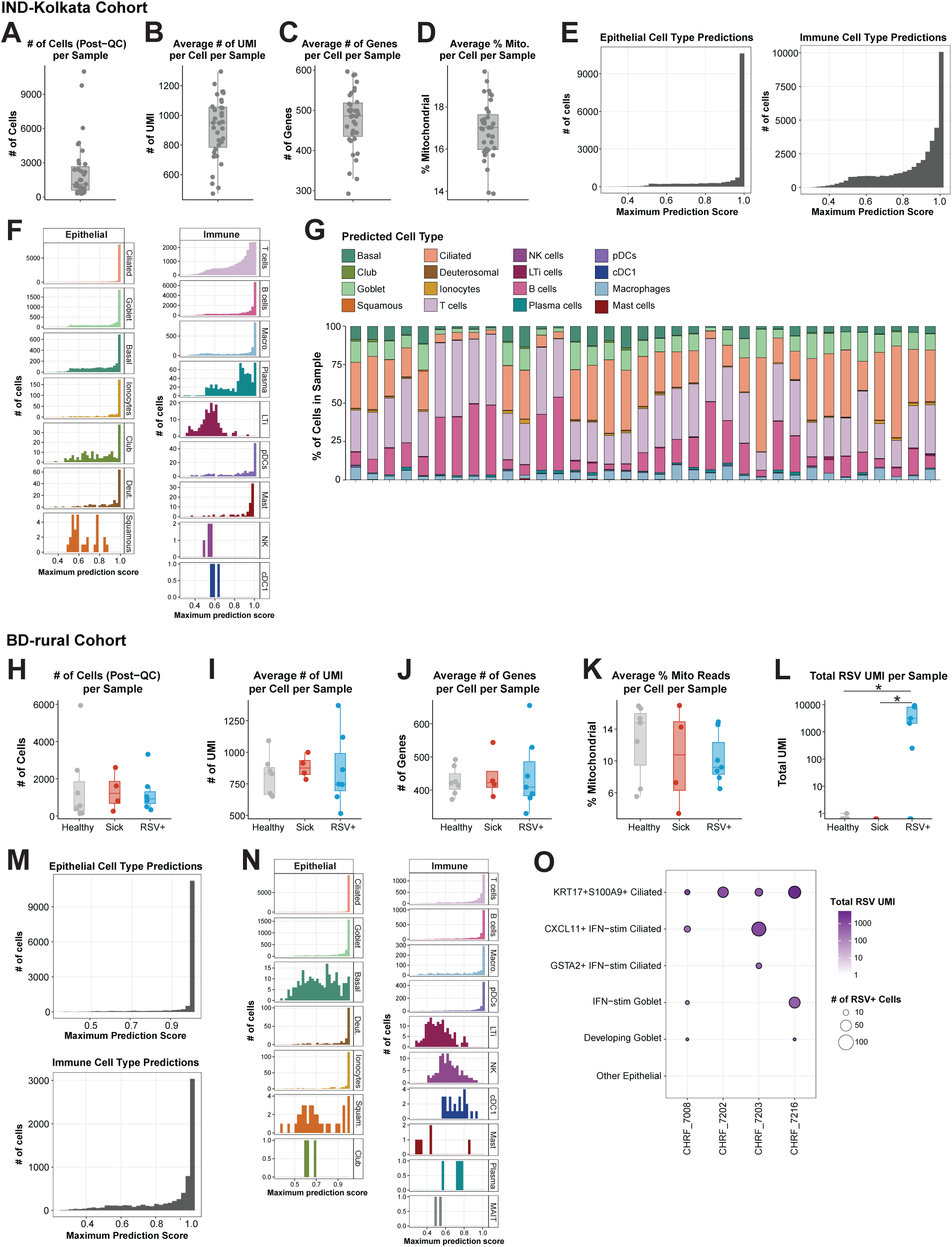
Related to Figure 7. **A-H.** Quality-control (QC) and prediction metrics for samples in IND-Kolkata cohort. **A.** Number of cells per sample after selecting high-quality cells (defined as >150 unique molecular identifiers (UMI), >100 unique genes, and <35% mitochondrial reads). Metrics in B-D are restricted to high-quality cells. **B.** Average number of UMI per cell per sample. **C.** Average number of unique genes per cell per sample. **D.** Average percent of mitochondrial reads per cell per sample. **E.** Distribution of maximum cell type prediction scores within all epithelial cells (left) and immune cells (right). **F.** Distribution of maximum cell type prediction scores divided by cell type. Cell types are colored as in **(G**). **G.** Distribution of predicted cell types as a percentage of all cells. Each column represents one sample. **H-O.** Quality-control and prediction metrics for samples in BD-rural cohort. **H.** Number of cells per sample after selecting high-quality cells (defined as >150 unique molecular identifiers (UMI), >100 unique genes, and <35% mitochondrial reads), divided by disease group. Metrics in I-K are restricted to high-quality cells. **I.** Average number of UMI per cell per sample by disease group. **J.** Average number of genes per cell per sample by disease group. **K.** Average percent of mitochondrial reads per cell per sample by disease group. **L.** Total number of RSV transcripts (UMI) per sample by disease group. **M.** Distribution of maximum cell type prediction scores within all epithelial cells (top) and immune cells (bottom). **N.** Distribution of maximum cell type prediction scores divided by cell type. Cell types are colored as in **(G**). **O.** Dotplot of RSV RNA+ cells in each epithelial subset, divided by sample. Samples with nonzero RSV RNA counts are shown. Dot size represents the number of cells that are viral RNA+. Color represents the total viral transcripts (UMI). **H-L:** Boxplots represent the median (center line), interquartile range (box), and 1.5x the interquartile range (whiskers). Statistical test is Kruskal-Wallis test with Benjamini-Hochberg correction for multiple comparisons. Dunn’s post-hoc test, *p<0.05, **p<0.001, ***p<0.0001.

## Supplementary Tables

**Supplementary Table 1.**
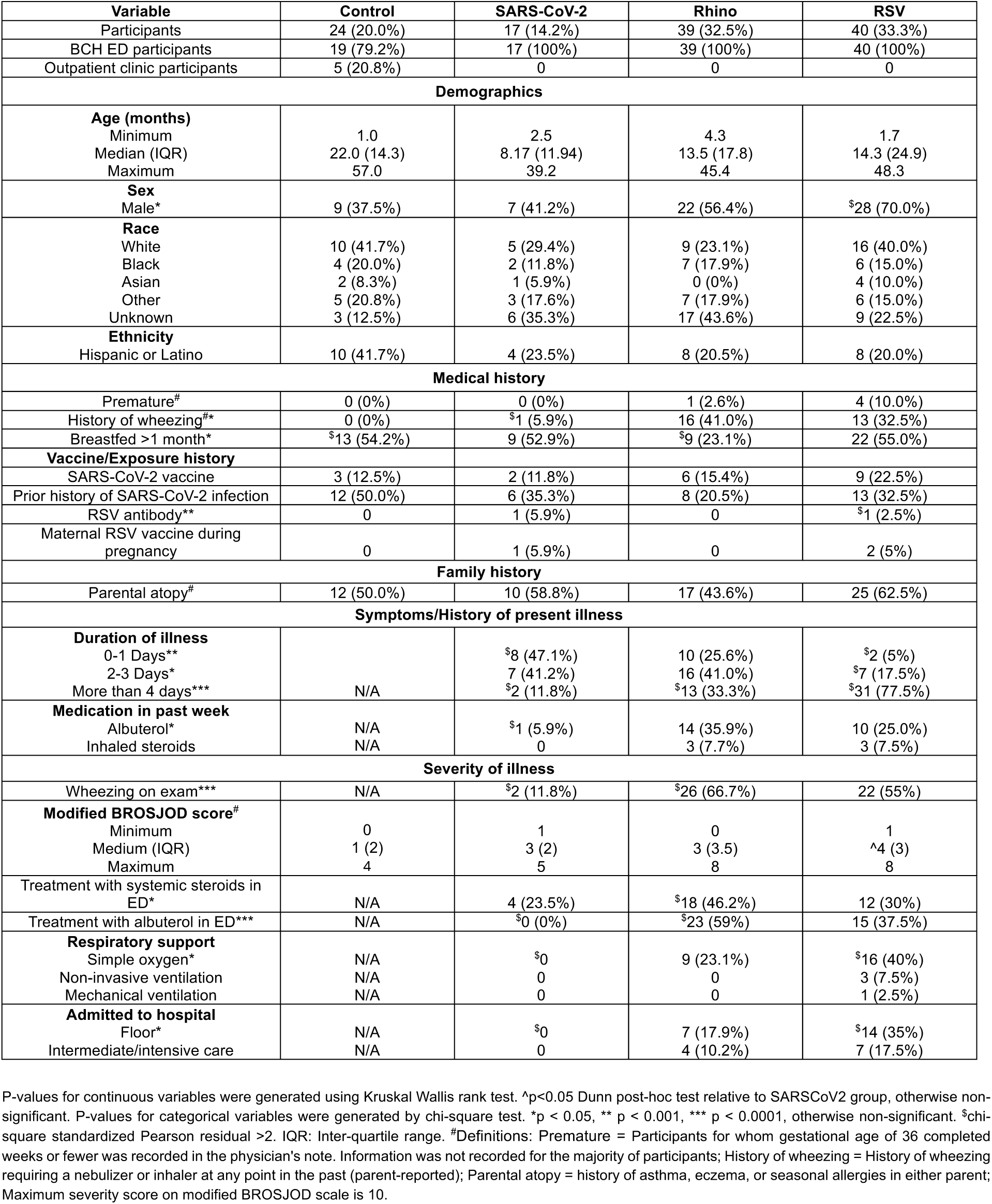
related to Figure 1: Summarized participant metadata for USA-Boston cohort.

**Supplementary Table 2.** related to Figures 1, 7 and Supplementary Figure 10: Full participant metadata for USA-Boston, IND-Kolkata, and BD-rural cohorts.

**Supplementary Table 3.** related to Supplementary Figures 2-3: Marker genes for final cell subsets defined within each cell type.

**Supplementary Table 4.** related to Figures 2F, 3B, and 6E: Differentially expressed genes displayed on volcano plots.

**Supplementary Table 5.** related to Figures 3E, 4D, 4E, 6F and Supplementary Figures 8F, 8G: Gene signatures used to calculate cytokine response scores.

**Supplementary Table 6.** related to Figure 5 and Supplementary Figure 6: Results from receptor-ligand analysis.

